# Variational and phase response analysis for limit cycles with hard boundaries, with applications to neuromechanical control problems

**DOI:** 10.1101/2022.05.17.492350

**Authors:** Yangyang Wang, Jeffrey P. Gill, Hillel J. Chiel, Peter J. Thomas

## Abstract

Motor systems show an overall robustness, but because they are highly nonlinear, understanding how they achieve robustness is difficult. In many rhythmic systems, robustness against perturbations involves response of both the shape and the timing of the trajectory. This makes the study of robustness even more challenging.

To understand how a motor system produces robust behaviors in a variable environment, we consider a neuromechanical model of motor patterns in the feeding apparatus of the marine mollusk *Aplysia californica* (Shaw et al., 2015; Lyttle et al., 2017). We established in (Wang et al., 2021) the tools for studying combined shape and timing responses of limit cycle systems under sustained perturbations and here apply them to study robustness of the neuromechanical model against increased mechanical load during swallowing. Interestingly, we discover that nonlinear biomechanical properties confer resilience by immediately increasing resistance to applied loads. In contrast, the effect of changed sensory feedback signal is significantly delayed by the firing rates’ hard boundary properties. Our analysis suggests that sensory feedback contributes to robustness in swallowing primarily by shifting the timing of neural activation involved in the power stroke of the motor cycle (retraction). This effect enables the system to generate stronger retractor muscle forces to compensate for the increased load, and hence achieve strong robustness.

The approaches that we are applying to understanding a neuromechanical model in *Aplysia*, and the results that we have obtained, are likely to provide insights into the function of other motor systems that encounter changing mechanical loads and hard boundaries, both due to mechanical and neuronal firing properties.

## 1 Introduction

In many animals, motor control involves neural oscillatory circuits that can produce rhythmic patterns of neural activity without receiving rhythmic inputs (central pattern generators (CPGs)), force generation by muscles, and interactions between the body and environment. Moreover, sensory feedback from the peripheral nervous system is known to modulate the rhythms of the electrical signals in CPGs and therefore facilitate adaptive behavior.

Motor systems show an overall robustness, but because they are highly nonlinear, understanding how they achieve robustness due to their different components is difficult. To understand how animals produce robust behavior in a variable environment, Shaw et al. (2015) and Lyttle et al. (2017) developed a neuromechanical model of triphasic motor patterns to describe the feeding behavior of the marine mollusk *Aplysia californica*. Like many rhythmic motor systems, feeding in *Aplysia* involves two distinct phases of movement: a *power stroke* during which the musculature engages with the substrate (the seaweed) against which it exerts a force to advance its goal (ingestion of seaweed), and a *recovery stage* during which the motor system disengages from the substrate to reposition itself, in preparation for beginning the next power stroke. Similarly, in legged locomotion, the stance phase corresponds to the power stroke and the swing phase corresponds to the recovery stage.

Also, like many rhythmic motor systems, feeding in *Aplysia* involves a closed-loop system, which integrates biomechanics and sensory feedback, and exhibits a stable limit cycle solution. It has been conjectured that sensory feedback plays a crucial role in creating robust behavior by extending or truncating specific phases of the motor pattern (Lyttle et al. (2017), §3.1). To test this hypothesis, we applied small mechanical perturbations as well as parametric perturbations to the sensory feedback pathways in the coupled neuromechanical system. It was shown in Lyttle et al. (2017) that a sustained increase in mechanical load leads to changes in both shape and timing of the limit cycle solution: the system generates stronger retractor muscle force for a longer time in response to the increased load. Qualitatively similar effects have been observed during *in vivo* experiments in *Aplysia* (Gill and Chiel, 2020). In general, we expect that applying parametric changes to CPG-based motor systems leads to changes in both the shape and timing of the resulting limit cycle behavior (Fig. 1).

**Fig. 1.**
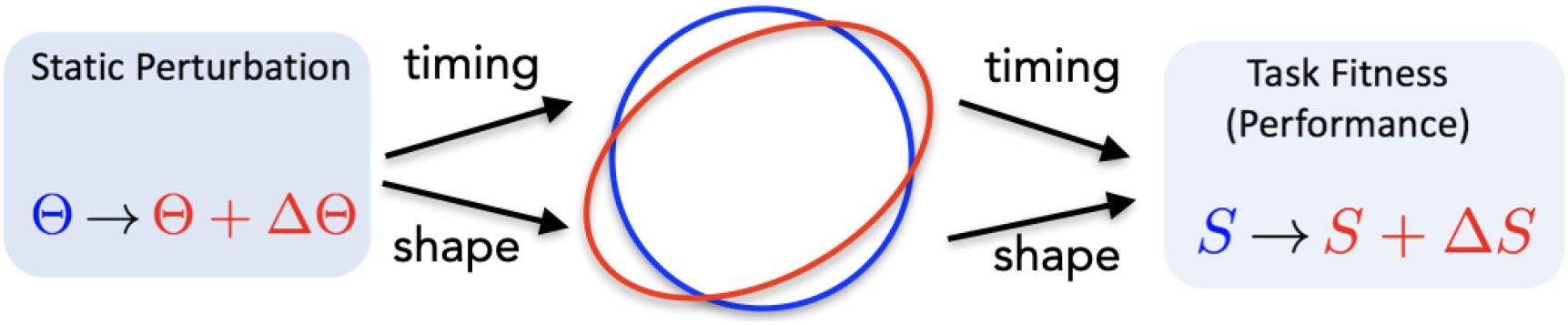
A sustained change in parameter *Θ* in a dynamical system 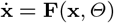 producing a limit cycle trajectory typically causes changes in both the timing and shape of the trajectory, which may both influence the performance *S* of the limit cycle system.

In *Aplysia*, the increased duration (timing) and increased force (shape) have opposite effects on the task-fitness, measured as seaweed consumption per unit time. Strengthening the retractor force pulls in more food with each cycle, which increases fitness, whereas lengthening the cycle time decreases fitness. Together these effects approximately cancel, making the system robust against increased load. This type of “stronger-and-longer” response may occur generically in other motor systems. Thus, in this paper, we seek to understand the roles of sensory feedback and biomechanics in enhancing robustness. To this end, we apply recently developed tools from variational analysis (Wang et al., 2021) to quantitatively study changes in shape and timing of a limit cycle under static perturbations.

In the first part of the present paper (cf. §2.1), we apply the classical tools of forward variational analysis to the model introduced by Shaw, Lyttle, Gill and coauthors in (Shaw et al., 2015; Lyttle et al., 2017) (denoted as the Shaw-Lyttle-Gill or SLG model for brevity) to arrive at the following insights:

– Nonlinear biomechanical properties confer resilience by immediately increasing resistance to applied loads, on timescales much faster than neural responses.
– The main effect of sensory feedback is to shift the timing of retraction neural pool deactivation; in parallel, firing rate saturation effectively censors sensory feedback during specific movement subintervals.

While the forward-in-time variational analysis is illuminating and allows us to explain in detail the robustness mechanism, it is still incomplete. Over time, the original and perturbed cycle will become increasingly out of phase due to the timing changes under sustained perturbations. Hence the shape displacements estimated from the forward variational analysis will become less and less accurate over time. This difficulty is not limited to models of feeding in *Aplysia californica*. For example, if we were to compare the gaits of two subjects walking on treadmills with slightly different speeds, although the ratio of stance and swing may be the functionally important aspect, this quantity is difficult to assess directly without putting the two movements on a common footing by comparing them using a common time scale.

Thus, in order to compare perturbed and unperturbed motions with greater accuracy, in the remainder of the paper (cf. §2.2 and following) we show how to extend the local-in-time variational analysis to a global analysis by applying the *infinitesimal shape response curve* (ISRC) analysis and *local timing response curve* (LTRC) analysis developed in Wang et al. (2021). We review these methods in §3. This time-rescaled analysis accounts for both global timing sensitivity (through the *infinitesimal phase response curve*, IPRC), as well as local timing sensitivity (through LTRC) by rescaling time to take into account local differences in the effects of parametric variation. It yields a more accurate and self-consistent description of the oscillator trajectory’s changing shape in response to parametric perturbations and helps complete the picture by providing a complementary perspective. Specifically, our time-rescaled analysis provides additional insights, specifically that

– Increasing the applied load on the system increases the duty cycle of the neuron pool responsible for retraction, in the sense that the retraction neuron pool is activated for a larger *percentage* of the closed phase of the cycle. (The *closed phase* of the trajectory occurs while the animal’s radula-odontophore, or grasper, is closed on the seaweed, and encompasses the power stroke.) This effect ultimately results in more seaweed being consumed, despite increased force opposing ingestion.
– We are able to derive the multidimensional *infinitesimal phase response curve* (IPRC) despite the presence of nonsmooth dynamics in the system; we identify the mechanical component of the IPRC as the one that contributes most to robustness, and note that its contribution arises from the “power stroke” segment of the motor cycle.
– We derive an analytical expression for the robustness to the mechanical perturbation that decomposes naturally into a sum of two terms, one capturing the effect of the perturbation on the shape of the trajectory, and the other capturing the effect on the timing; this result provides a *quantitative* analysis of robustness that confirms the *qualitative* insights described previously in the literature.
– In addition to sensory feedback and intrinsic biomechanical properties, robustness against changes in applied load can arise from coordinated changes of multiple parameters such as the gain of sensory feedback and muscle stiffness.

The dynamics of the SLG model (Lyttle et al., 2017) is given by

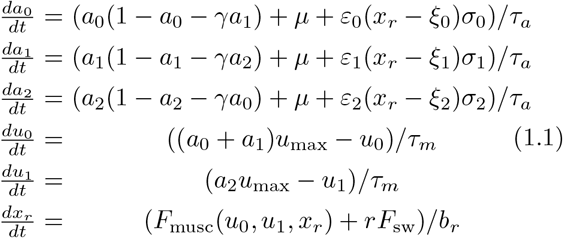

This system incorporates firing rates of three neuron populations, corresponding to the “protraction-open” (*a*_0_), “protraction-closed” (*a*_1_), and “retraction” phase (*a*_2_). Note that when a nerve cell ceases firing because of inhibition, its firing rate will be held at zero until the balance of inhibition and excitation allow firing activity to resume. Hence, we supplement model (1.1) with three hard boundaries introduced by the requirement that the firing rates *a*_0_, *a*_1_, *a*_2_ must be nonnegative:

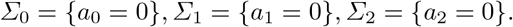

During the limit cycle, when a neural variable *a*_*i*_ changes from positive to 0, we call that the *a*_*i*_ *landing point*; when it changes from 0 to positive, we call that the *a*_*i*_ *liftoff point*. The fact that the trajectory is non-smooth at the landing and liftoff points will play an important role in the analysis to follow.

This model also consists of a simplified version of the mechanics of the feeding apparatus: a grasper that can open or close (*x*_*r*_), a muscle that can protract the grasper to reach the food (*u*_0_) and another muscle that can retract the grasper to pull the food back into its mouth (*u*_1_). The net force exerted by the muscles is given by the sum of the two muscle forces

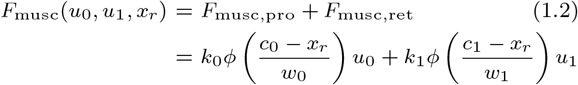

where

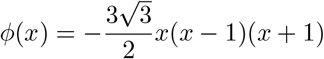

is the effective length-tension curve for muscle forces, *c*_*i*_, *w*_*i*_ and *k*_*i*_ denote the mechanical properties of each muscle.

*F*_sw_ represents the external force applied to the seaweed, which can only be felt by the grasper when it is closed on the food (*a*_1_ + *a*_2_ *>* 0.5), during which *r* = 1. When the grasper is open (*a*_1_ + *a*_2_ ≤ 0.5), *r* = 0; that is, the grasper moves independently of the seaweed. This condition leads to a transversal crossing boundary; that is, the open/closing boundary given by

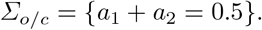

Values for model parameters and initial conditions are given in Table 2 and Table 3 in the Appendix. For additional details on the biological assumptions motivating the model, see (Shaw et al., 2015; Lyttle et al., 2017).

**Table 2.** Model parameters

| Parameter | Value | Description |
| --- | --- | --- |
| $\gamma$ | 2.4 | inhibition strength from next pool |
| $\varepsilon_i$ | $10^{-4}$ | sensory feedback strength |
| $\mu$ | $10^{-6}$ | neural pool intrinsic excitation |
| $\tau_a$ | 0.05 | neural pool time constant |
| $\tau_m$ | 2.45 | muscle activation time constant |
| $b_r$ | 0.4 | grasper damping constant |
| $c_0$ | 1.0 | position of shortest length for I2 |
| $c_1$ | 1.1 | position of center of I3 |
| $F_{sw}$ | 0.01 | force on the seaweed resisting ingestion |
| $\sigma_0$ | -1 | sign of proprioceptive input to $a_0$ motor pool |
| $\sigma_1$ | 1 | sign of proprioceptive input to $a_1$ motor pool |
| $\sigma_2$ | 1 | sign of proprioceptive input to $a_2$ motor pool |
| $\xi_0$ | 0.5 | proprioceptive neutral position for protraction-open neural pool |
| $\xi_1$ | 0.5 | proprioceptive neutral position for protraction-closed neural pool |
| $\xi_2$ | 0.25 | proprioceptive neutral position for retraction-closed neural pool |
| $u_{max}$ | 1.0 | maximum muscle activation |
| $w_0$ | 2 | maximal effective length of I2 |
| $w_1$ | 1.1 | maximal effective length of I3 |
| $k_0$ | 1 | strength and direction of the protractor muscle |
| $k_1$ | -1 | strength and direction of the retractor muscle |

**Table 3.** State variables

| State variable | Initial value | Description |
| --- | --- | --- |
| $a_0$ | 0.9 | activity of I2 motor pool (non-negative) |
| $a_1$ | 0.08355 | activity of hinge motor pool (non-negative) |
| $a_2$ | 0.00003 | activity of I3 motor pool (non-negative) |
| $u_0$ | 0.748 | activity of I2 muscle |
| $u_1$ | 0.25 | activity of I3 muscle |
| $x_r$ | 0.65 | grasper position (0 is retracted, 1 is protracted) |

In this paper, we are interested in the so-called heteroclinic mode in which the neural dynamics temporarily slow down when the sensory feedback overcomes the endogeneous neural excitation and forces the neural trajectory to slide along the hard boundary *a*_*i*_ = 0. Such temporary slowing down of neural variables allows the muscles, which evolve on slower timescales, to “catch up”; hence the seaweed can be swallowed and ingested successfully. Following the terminology from (Wang et al., 2021), we call attracting periodic trajectories that experience sliding motions *limit cycles with sliding components* (LCSCs).

Using classical sensitivity analysis and our recently developed tools from variational analysis (Wang et al., 2021), we show here that biomechanics and sensory feedback cooperatively support strong robustness by changing the timing and shape of the neuromechanical trajectory. While both sensory feedback and biomechanics respond immediately to the increased load, we find that the sensory feedback effect is initially censored while the neural activity is pinned against a hard boundary of neuronal firing. Thus the effect of the sensory feedback signal is significantly delayed relative to onset of the increased load. Our analysis suggests that sensory feedback mediates robustness mainly through shifting the timing of neural activation and specifically increasing the duty cycle of the retraction neural pool. This response allows the system to digest more seaweed despite the increased force opposing ingestion and hence achieve strong robustness. In addition to uncovering the mechanisms for robust motor control, our methods allow us to quantify analytically the robustness of the model system to the mechanical perturbation. Finally, although we focus on the *Aplysia californica* feeding system as our working example, our methods should extend naturally to a broad range of motor systems.

Our paper is organized as follows. We present our analysis and main results in §2. Methods that we use to understand the robustness in the *Aplysia* neuromechanical model are presented and reviewed in §3. We discuss limitations and possible extensions of our approach in §4.

## 2 Results

Recall that a sustained (parametric) perturbation often causes changes in both shape and timing of the neuromechanical trajectory solution of (1.1). In this paper, we adopt methods developed in (Wang et al., 2021) for analyzing the joint variation of both shape and timing of limit cycles with sliding components under parametric perturbations.

### 2.1 Forward Variational Analysis

We begin our analysis by investigating how the shape of the trajectory changes in response to a sustained increase in mechanical load *F*_sw_. To a first approximation, the change in shape can be captured by classical sensitivity analysis (also called forward variational analysis) which we review in §3.

We apply a small static perturbation to the system (1.1) by increasing the model parameter *F*_sw_ by *ε* = 0.02: *F*_sw_ → *F*_sw_ + *ε*, and comparing the new, perturbed limit cycle trajectory *γ*_*ε*_ to the original, unperturbed limit cycle trajectory *γ*_0_, beginning at the start of the grasper-closed phase (time 0 in Figure 2 panels A-D). That is, we plot **u** ≈ (*γ*_*ε*_(*t*) − *γ*_0_(*t*))*/ε*; see (3.13)-(3.15) for precise definitions. Note that *F*_sw_ is multiplied by an indicator function that is only nonzero when the trajectory is in the grasper-closed phase. Thus the perturbation is only present when the grasper is closed on the food.

The neural and biomechanical components of the unperturbed trajectory *γ*(*t*) are shown by the solid curves in Figure 2A and B, respectively. The perturbed trajectories *γ*_*ε*_(*t*) are indicated by the dashed lines. The gray shaded regions indicate phases when the grasper in the unperturbed system is closed. With the perturbation (increased load), the transition from closing to opening is delayed; this transition is indicated by the magenta vertical line. Figure 2C and D show the difference between the two trajectories per perturbation along the neural directions and along the biomechanical directions, respectively. These curves can be approximated by the solutions to the forward variational equation (3.14) defined in §3. The muscle forces *F*_musc_(*u*_0_, *u*_1_, *x*_*r*_) before and after the perturbation are shown by the gray curves in Figure 2B), and the difference between them is included as the gray curve in Figure 2D).

**Fig. 2.**
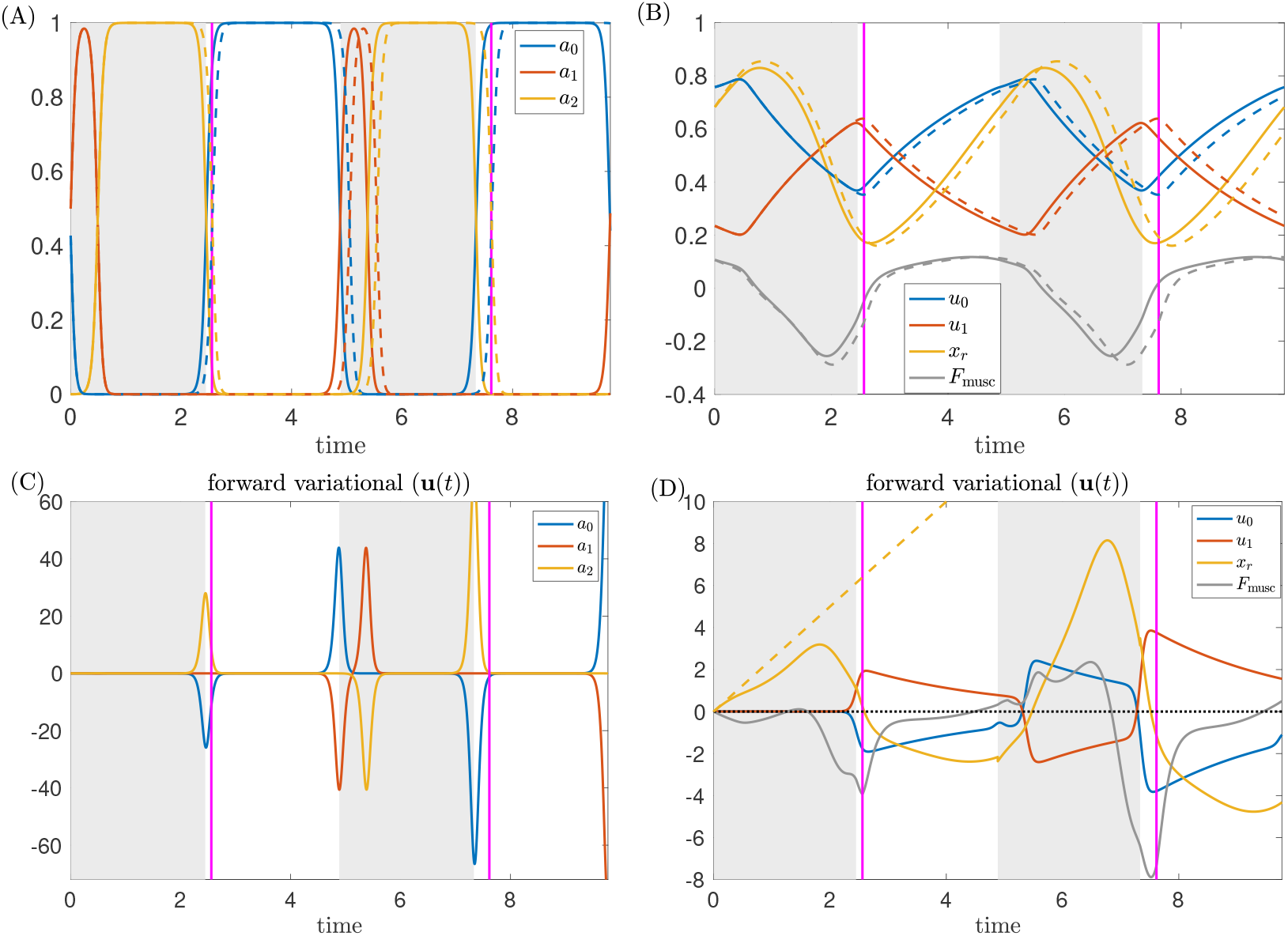
A small sustained perturbation is applied to the *Aplysia* model (1.1) over the closing phase in which *F*_sw_ → *F*_sw_ +*ε* with perturbation *ε* = 0.02. (A, B) Time series of the trajectory components for nominal force value *F*_sw_ (solid) and perturbed force value (dashed) over two periods, aligned at the start of the closed phase. (C, D) The displacement solution, **u**(*t*), to the forward variational equation over two periods. The gray curve in (D) denotes the displacement between the perturbed and unperturbed muscle forces *F*_musc_, shown as the gray curves in panel (B). The yellow dashed line in panel (D) approximates the displacement in *x*_*r*_ if the net muscle force *F*_musc_ did not change after perturbation. The intervals during which the grasper without perturbation is closed on the food are indicated by the shaded regions. The vertical magenta lines indicate the times at which the grasper under perturbation switches from closed to open. The difference in periods and the delay in the grasper opening time both accumulate, making the comparison between the two trajectories invalid except for short times. (A) and (C) show trajectories and displacements along the neural directions, while (B) and (D) show trajectories and displacements along the mechanical directions. The lines in panel (C) (resp., (D)) approximate the difference between the dashed and solid trajectories from panel (A) (resp., (B)) per perturbation.

Figure 2 yields several insights about the roles of sensory feedback and biomechanics in robustness, which we discuss in detail below.

#### Biomechanics confer resilience by immediately increasing resistance to the increased load

Immediately following the perturbation of the mechanical load, we observed a positive displacement in the grasper position *x*_*r*_ relative to the unperturbed trajectory (Figure 2D, yellow curve). This displacement simply reflects the grasper being pulled by a stronger force *F*_sw_ + *ε*. If nothing other than the applied load *F*_sw_ changes in the system, a linearized analysis suggests that the initial displacement of *x*_*r*_ would approximately follow the yellow dashed line given by 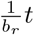 (Figure 2D). Nonetheless, while this line gives a good initial approximation, the true displacement in *x*_*r*_ (yellow solid curve) quickly sags below the yellow dashed line over time. This difference arises due to the negative displacement occurring in the muscle force *F*_musc_(*u*_0_, *u*_1_, *x*_*r*_) (Figure 2D, gray curve) which acts to overcome the increased load. However, early in the retraction cycle, all other variables, including the muscle activation *u*_0_ and *u*_1_, show almost no displacement at all (see Figure 2C and D). This observation suggests that long before the sensory feedback effect has time to act, the biomechanics may play an essential role in generating robust motor behavior, by providing an immediate, short-term resistance to increased load.

Early in the retraction cycle, increasing the load stretches both the retractor and protractor muscles, and moves them down their length-tension curves. As a result, both forces become weaker, but the magnitude of the protractor muscle force drops more quickly than the retractor muscle force (Figure 3). Thus, the retractor muscle force grows relative to the opposing protractor muscle force. This shift endows the system with a built-in resilience, in that increasing seaweed force opposing ingestion automatically (i.e., without changes in neural activation) engages a larger resisting muscle force long before neural pools or muscles show differential activation. This is a new insight beyond the previous “longer-stronger” hypothesis (Shaw et al., 2015; Lyttle et al., 2017).

**Fig. 3.**
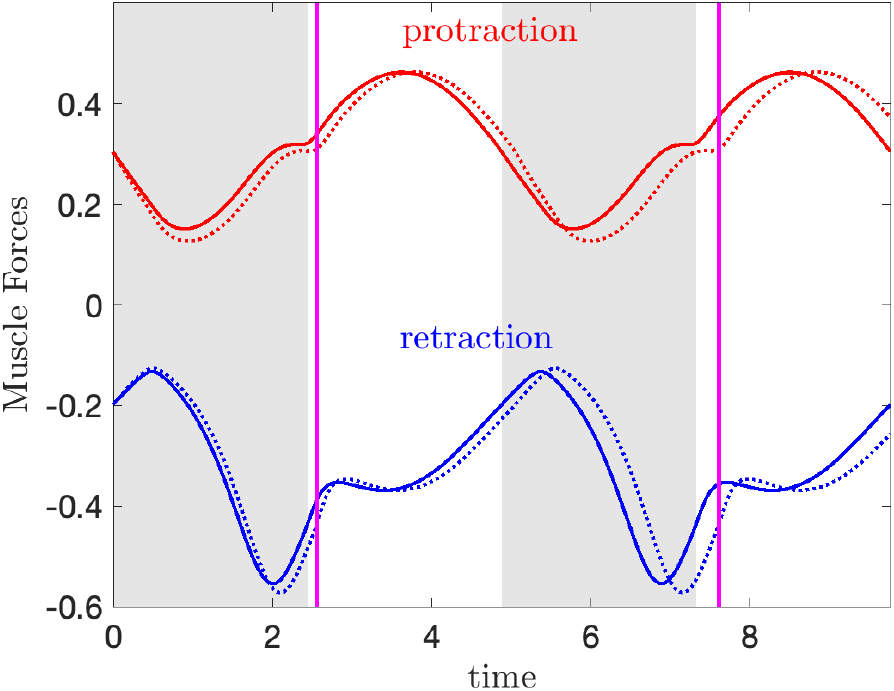
The time series of the perturbed (dashed) and unperturbed (solid) protractor muscle forces *F*_musc,pro_ (red) and retractor muscle forces *F*_musc,ret_ (blue) over two periods. The gray shaded regions and the magenta lines have the same meanings as in Figure 2.

#### Sensory feedback effects are largely delayed by the firing rate hard boundary properties

Changes in *x*_*r*_ due to the increased load will immediately propagate to the neural variables (*a*_0_, *a*_1_, *a*_2_) through the sensory feedback *ε*_*i*_(*x*_*r*_ − *ξ*_*i*_)*σ*_*i*_)*/τ*_*a*_, and hence should affect the neural variables without any lag. However, no significant displacement of the neural variables is observed until nearly the end of the retraction cycle (Figure 2C). In other words, while the sensory feedback itself immediately responds to the increased load, the *effect* of the changed sensory feedback signal is not manifest until much later in the retraction cycle, when the protraction-open neuron pool is released from inhibition along its hard boundary and starts to fire (Figure 2). Hence, the nonsmooth hard boundary conditions on neuronal firing rates significantly delay the effect of sensory feedback, and create intervals during which neurons are insensitive to sensory feedback. This effect is a concrete example of *differential penetrance* (Ye et al., 2006; Cullins et al., 2015).

#### *Sensory feedback contributes by shifting the* timing *of neural activation*

Due to the hard boundary effects, the displacements of the neural variables appear near the end of the closed phase, when the protraction-open neuron pool *a*_0_ lifts off from its hard boundary, and the retraction-closed neuron pool *a*_2_ deactivates to stop firing (Figure 2A and C). A positive (resp., negative) displacement of *a*_2_ (resp., *a*_0_) indicates that *a*_2_ deactivates (resp., *a*_0_ activates) at a later time with the increased load, and hence the retraction-closed phase is prolonged. Consequently, the retraction muscle activity will increase, because its stimulation by the retraction motor neuron is prolonged, allowing the slow retractor muscle to generate larger forces (Figure 3D). Similarly, we also observe a decreased protractor muscle activity, as the protraction neuron pool *a*_0_ turns on at a later time. This *decrease* leads to a stronger retractor muscle force and a weaker protractor muscle force (Figure 3). Hence, a more negative net muscle force results (Figure 2D, gray curve), which corresponds to a stronger resisting force pulling the seaweed towards the jaw to swallow the food. To summarize, the main effect of sensory feedback that contributes to robustness is prolonging the retraction phase to confer on the system a resilience in that increasing seaweed force opposing ingestion engages a larger resisting muscle force. Thus, sensory feedback contributes to robustness primarily by shifting the *timing* of neural activation, as opposed to the magnitude of neural activation. Biologically, this distinction corresponds to affecting the timing of motor neuron burst onset or offset, rather than burst intensity.

### 2.2 Variational Analysis with Rescaled Time - ISRC

Under the forward-in-time analysis, the grasper of the perturbed system lags behind the grasper of the unperturbed system throughout the closing phase; yet the *net* seaweed movement was measured to be greater for the perturbed system (Lyttle et al., 2017). Fig. 4 shows an expanded view of the perturbed and unperturbed systems’ grasper position (Fig. 4B) and the linearized difference produced by the variational equation (Fig. 4D). As this detailed view shows, at the time when the unperturbed system transitions from closed to open (gray-white boundary) the unperturbed grasper position is *more retracted* than the perturbed grasper position at the coincident time point. Similarly, the grasper component of the variational equation is *positive* at the gray-white boundary. Furthermore, at the time when the perturbed system transitions from closed to open (magenta line) the perturbed system continues to be less retracted than the unperturbed system. Thus, whether we compare the systems at the perturbed or unperturbed opening time, the perturbed grasper is “further behind”. Yet the overall effect in the perturbed system is a larger net intake of seaweed per cycle.

**Fig. 4.**
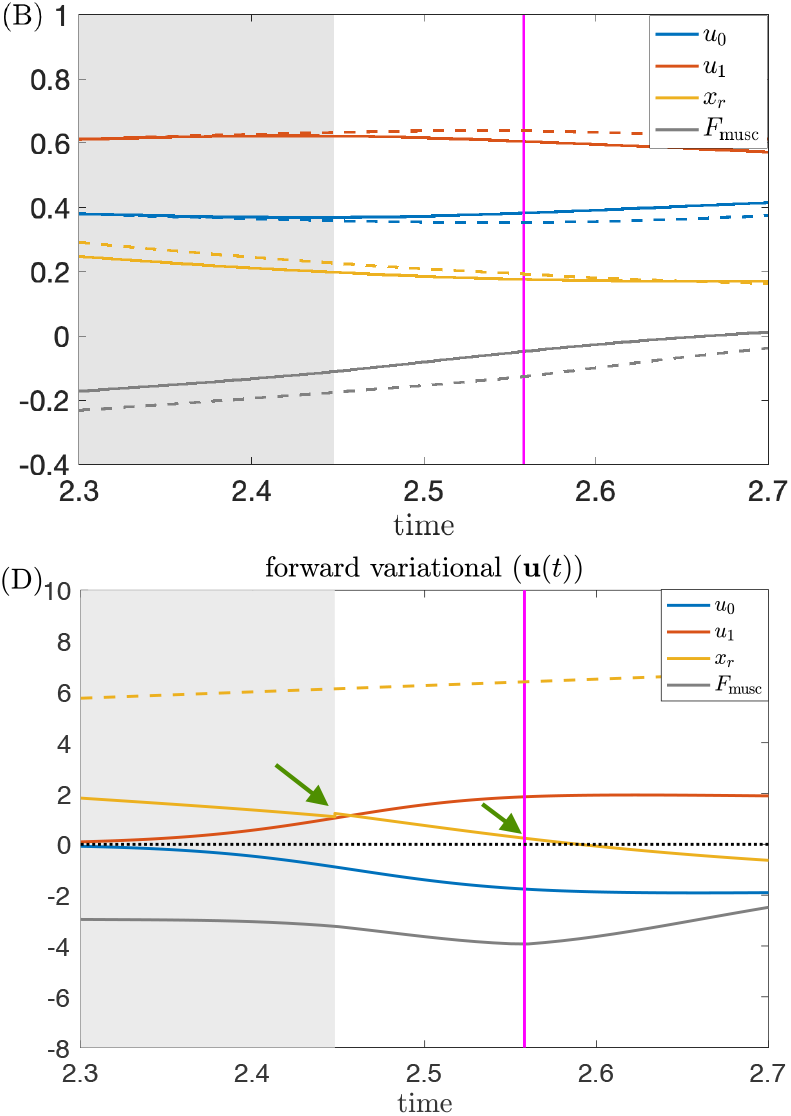
Enlarged views of Figure 2B and D near the first transition from closed to open. Note the displacement of *x*_*r*_ (i.e., the *x*_*r*_ component of **u**(*t*) shown as the yellow solid line in (D)) is positive both at the closing time of the unperturbed system (see the green arrow near the grey/white boundary) and the perturbed system (see the green arrow near the magenta line).

This apparent contradiction underscores the need to extend the perturbation analysis beyond the standard forward-in-time variational analysis. In particular, if one cycle is slower than another, then while the local perturbation analysis can explain the cause-and-effect relations a short time into the future, they cannot account for the net effect around a cycle in a self-consistent way. Over time, the displacements between the two trajectories grow, and the linearized approximation becomes invalid except at short times (cf. Jordan et al. (2007)). Hence, unless time is rescaled to take into account the difference in cycle period, comparing the components of the original and perturbed cycles will become less and less meaningful.

To overcome this difficulty, we extend the local-in-time variational analysis to a global analysis by rescaling time so the unperturbed closing and opening events coincide with those after perturbations, respectively. We do so by applying the *infinitesimal shape response curve* (ISRC) analysis and the *local timing response curve* (LTRC) (Wang et al., 2021), which we review in §3. This method yields a more accurate and self-consistent description of the oscillator trajectory’s changing shape in response to parametric perturbations (see Figure 5). We show that the combination of the ISRC and the LTRC gives a sensitivity analysis of an oscillator to sustained perturbations within any given region (e.g., protraction or retraction cycle, opening or closing phase) and provides a self-contained framework for analytically quantifying and understanding robustness to perturbations.

**Fig. 5.**
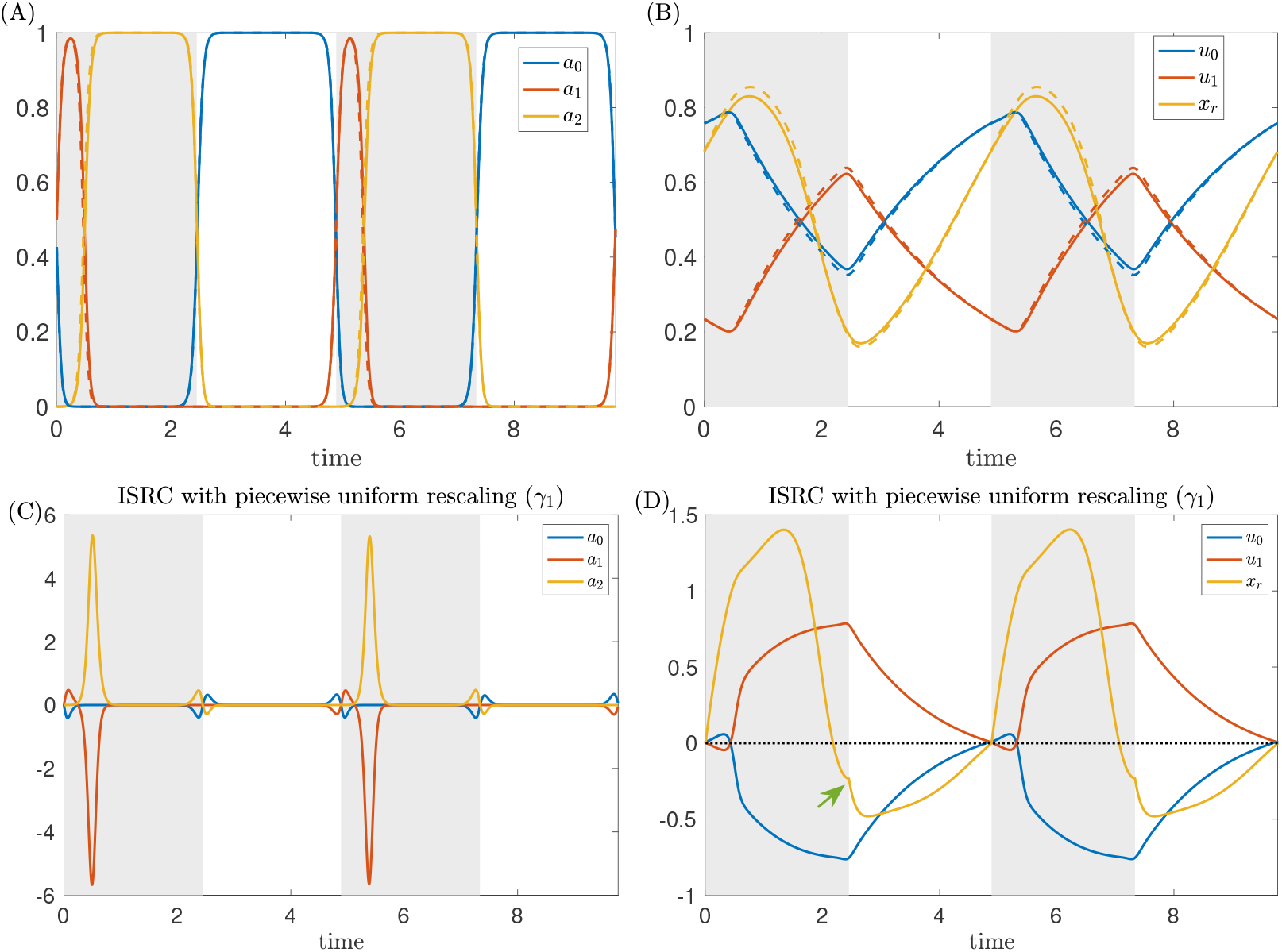
Variational analysis with piecewise uniform time rescaling. The same sustained perturbation as in Figure 2 is applied to the *Aplysia* model (1.1). (A, B) Time series of the perturbed (dashed) and unperturbed solutions (solid). Here piecewise uniform rescaling is applied so the closing and opening events coincide. (C, D) The ISRC with piecewise uniform rescaling *γ*(*t*) over two periods. Shaded regions have the same meanings as in Figure 2. Note the *x*_*r*_ component of the ISRC is negative at the time of opening (see green arrow). With piecewise uniform rescaling, the variational approximation is consistent across multiple periods (c.f., Figure 2).

We write *γ*_1_ for the linear shift in the limit cycle shape in response to the static perturbation *F*_sw_ → *F*_sw_ + *ε*, that is:

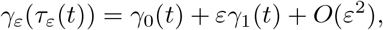

uniformly in time. Note that the time for the perturbed trajectory is rescaled to be *τ*_*ε*_(*t*) to match the unperturbed time points. The linear shift *γ*_1_(*t*) is the so-called ISRC curve and satisfies a nonhomogeneous variational equation (see §3). Compared with the forward variational equation, the ISRC equation has one additional nonhomogeneous term *ν*_1_*F*_0_(*γ*_0_(*t*)) that arises from the time rescaling. In this term, *ν*_1_ is determined by the choice of time rescaling *τ*_*ε*_(*t*) and *F*_0_(*γ*_0_(*t*)) is the unperturbed vector field evaluated along the unperturbed limit cycle *γ*_0_(*t*) (see §3 for details).

Since the perturbation is applied to the seaweed, it can only be felt by the system when the grasper is closed on the seaweed. It is natural to expect that the segment at the closing phase has a different timing sensitivity than the segment at the opening phase. We hence choose to rescale time differently in the two phases, using piecewise uniform rescaling when computing the ISRC. This leads to a piecewise ISRC equation, where *ν*_1_ is piecewise constant. It was shown in (Wang et al., 2021) that *ν*_1_ can be estimated from the LTRC analysis (see §3).

In Figure 5A and B, the time traces of variables along the unperturbed limit cycle are shown by the solid curves, whereas the perturbed limit cycle whose time has been rescaled to match the unperturbed time points as described above are indicated by the dashed curves. With the piecewise rescaling, the transitions between the closing and opening events of the perturbed and unperturbed systems now coincide. The relative displacements between the perturbed and unperturbed trajectories are approximately given by the piecewise ISRC *γ*_1_ shown in Figure 5C and D. In contrast to the forward variational analysis, in which the displacements grow over time, the piecewise ISRC curve is periodic, meaning we have achieved a self-consistent global description of the response of the limit cycle to increased load.

We now show that the apparent contradiction that we obtained from the forward variational analysis, i.e., that the grasper displacement at the end of the closing phase is positive (cf. Fig.4), can now be resolved in the time-rescaled picture. In response to the perturbation, the relative displacement of the grasper position (the *x*_*r*_ component of *γ*_1_, detoted as 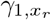) initially increases (i.e., the grasper becomes more and more protracted due to the increased load) and reaches its peak at about *t* = 1.4 (see Figure 5D, yellow curve). Then it starts to decrease and becomes negative at the time when the grasper opens. This means later in the retraction cycle, the perturbed grasper is less and less protracted than the unperturbed version and eventually become more retracted by the end of the closing phase (Fig. 5D, green arrow). In summary, the grasper perturbed by larger force begins “behind” the unperturbed version, but catches up around 60% of the way through the retraction phase (in relative time) and comes out “ahead” by the time both graspers open, consistent with having a larger net seaweed intake (Lyttle et al., 2017).

To understand what causes 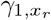 to be negative despite its initial big rise, we consider the effect of the perturbation on the neural pool through sensory feedback. In Figure 5C, we observe positive displacements in 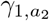 (yellow curve) occurring both when the retraction neuron pool *a*_2_ activates and when it deactivates. These displacements indicate that with the increased load, the retraction neuron *a*_2_ activates earlier and turns off later relative to the unperturbed *a*_2_. In other words, increasing the applied load on the system increases the *duty cycle* of the neuron pool involved in retraction, i.e., the retraction neuron pool is activated for a larger percentage of the total cycle. As a result, the motor system recruits a larger retractor muscle force, as indicated by the positive displacement of the retractor muscle activation *u*_1_ during the closing phase (Figure 5D, red curve). A similar increase in motor recruitment in response to increased external load has been observed *in vivo* (Gill and Chiel, 2020). In the model, the stronger retraction force acts to impede the protraction of the grasper, and eventually pulls the grasper to a more retracted state. Thus the grasper displacement crosses zero and becomes negative at the end of the closing phase (Figure 5D, green arrow).

Note that there is no perturbation during the opening phase (Figure 5, white space). During this phase of the cycle, displacements slowly decay and are nearly zero by the time the grasper closes on the food again.

### 2.3 Timing responses to sustained perturbations of *F*_sw_

#### Infinitesimal phase response curve

To understand the timing response of system (1.1) to increased load, we perform an IPRC analysis. Figure 6 shows the time traces of the IPRC curve over one cycle. As before, the shaded region indicates the phase when the grasper is closed.

**Fig. 6.**
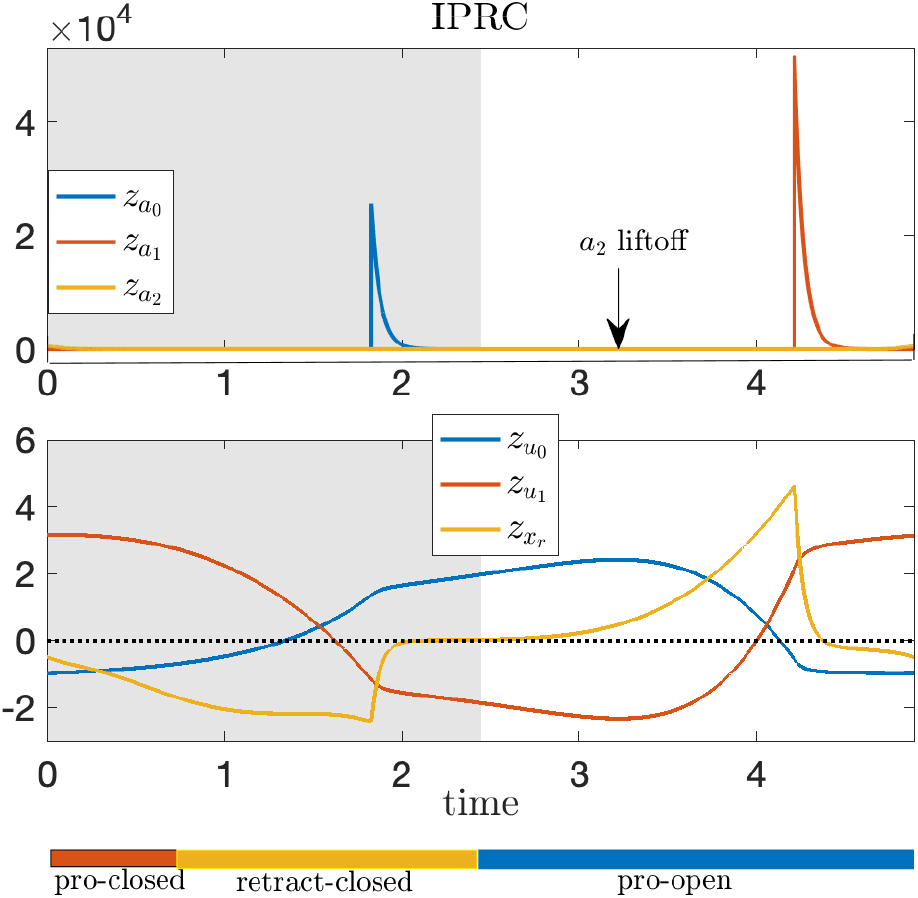
IPRC for the *Aplysia* model. Grey shaded region indicates the period when the radula/odontophore is closed. On the bottom, the red, yellow and blue rectangles denote the protraction-closed, retraction-closed and protraction-open phases, respectively. The blue spike in the IPRC in the top panel occurs when the *a*_0_ variable “lifts off” from the *a*_0_ = 0 boundary; the red spike occurs when the *a*_1_ variable lifts off; the liftoff point for *a*_2_ is indicated with an arrow.

The IPRC curves associated with biomechanical variables are shown in Figure 6, lower panel. In particular, the timing sensitivity of system (1.1) to the increased load on the grasper (*F*_sw_ → *F*_sw_ + *ε*) can be estimated using the IPRC along the *x*_*r*_ direction, i.e., the yellow curve in the lower panel of Figure 6. Since the perturbation only has effect during the closing phase, only the portion of 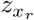 over the shaded region is relevant. This portion is strictly negative. Therefore, in response to the increased load considered above, the system undergoes phase delay, and hence the total period is prolonged. This finding is consistent with earlier results on the sensory feedback effect obtained from the variational analysis (see §2.2).

The linear shift in period can be estimated by evaluating the integral

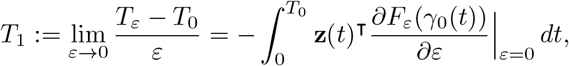

where *T*_0_, *T*_*ε*_ are the periods before and after perturbation *ε* (see Section 3). For the perturbation on *F*_sw_, the derivative 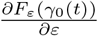 equals 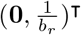 over the grasper-closed region, and equals **0** during the grasper-open region, where the first **0** is a 5 *×* 1 zero vector and the second **0** is a 6 *×* 1 vector. It then follows that

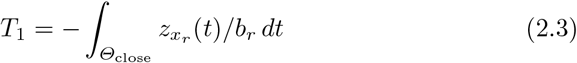

where *Θ*_close_ denotes the grasper-closed phase.

Other IPRC curves in Figure 6 indicate the timing sensitivity of the model to other perturbations and lead to several useful insights as well as testable predictions. For example,

– The IPRC curves are continuous except at the liftoff points (Figure 6 top panel, blue and red spikes). While all three neural variables go through liftoff points, there is no large spike in 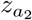 (yellow curve). The absence of a yellow spike and the fact that the red spike is larger than the blue spike, imply that the system has the highest timing sensitivity to perturbing *a*_1_ and intermediate timing sensitivity to *a*_0_, both of which are significantly higher than the sensitivity to *a*_2_ perturbations.
– Excitatory inputs to neural populations lead to phase advance and hence shorten the total period, because the IPRC curves associated with neural variables are mostly positive (Figure 6 top panel).
– Most of the time the system is not sensitive to neural perturbations, but there also exist sensitive regions when the trajectory is not restricted to the hard boundaries (e.g., Figure 6 top panel, blue and red spikes). For instance, the system has high timing sensitivity to perturbations of *a*_0_ late in the closing phase and to perturbations of *a*_1_ late in the opening phase, whereas sensory inputs are largely ignored early in the opening phase.
– Increasing the protractor muscle activation *u*_0_ causes a phase delay early in the closing phase and late in the opening phase, and a phase advance otherwise. In contrast, increasing the retraction muscle activation *u*_1_ causes a phase advance early in the closing phase and late in the opening phase, and a phase delay otherwise. Appendix B discusses why the system has different timing sensitivities to muscle perturbations.

Although all three neural variables go through liftoff points, there is no large yellow spike in 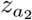 (see Figure 6). To understand this, we note that before *a*_0_ (resp., *a*_1_) lifts off its hard boundary, there exists no inhibition from other neurons except for inhibitory sensory feedback. However, when *a*_2_ lifts off at around *t* ≈ 3.2, it still experiences inhibition from *a*_0_ (see Figure 5A). In other words, there are two inhibitory effects pressing neurons *a*_2_ down to the hard boundary, but only one inhibitory effect acting on the other two neuron populations. As a result, while there is a discontinuous jump of the IPRC curve corresponding to *a*_2_ at the liftoff point, it remains small as the other inhibition is still present.

#### Local timing response curve

While the IPRC is a powerful tool for understanding the global timing sensitivity of an oscillator to sustained perturbations, it does not give local timing sensitivities, which, however, are needed for computing the ISRC curve as discussed above. We hence adopt the local timing response curve (LTRC) method developed in Wang et al. (2021) and reviewed in §3. To illustrate this method, we show the LTRC associated with the closing phase and denote it as *η*^close^ (see Figure 7). Although the LTRC *η*^close^ is defined throughout the full domain, estimating the effect of the perturbation within the closing region only requires evaluating the LTRC in this region. Figure 7 shows the time series of *η*^close^ for the model in the closing region, obtained by numerically integrating the adjoint equation backward in time with the initial condition of *η*^close^ given by its value when the grasper switches from closing to opening. Note that 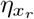, the yellow curve in Figure 7 lower panel, remains positive over the closing phase. This implies that the increased load on seaweed prolongs the time remaining in the closing region; that is, the increased load prolongs the total closing time. The relative shift in the closing time caused by the increased load can also be estimated by integrating the LTRC (see Section 3).

**Fig. 7.**
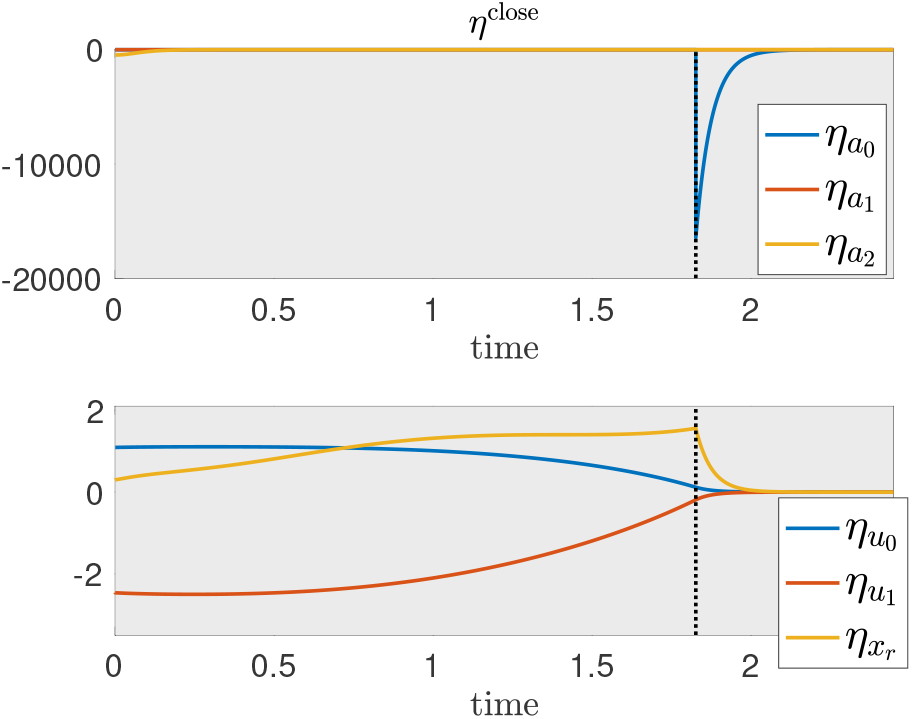
Time series of the LTRC *η*^close^ over the closing phase. The liftoff point on *a*_0_ = 0 coincides with the spike in 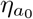 (blue curve, top panel). The cusp where 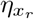 changes from increasing to decreasing (intersection of yellow and vertical black dashed curves in the bottom panel) also occurs at the liftoff point for *a*_0_.

In addition, Figure 7 implies that strengthening the protractor muscle activation *u*_0_ during the closing phase prolongs the total closing time, whereas increasing the retraction muscle activation *u*_1_ decreases the total closing time. Similarly, we can compute the LTRC over other phases, such as the retraction phase, in order to estimate local timing sensitivities of the system in other regions.

Finally we note an interesting feature in *η*^close^: there is an abrupt change in 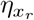 at the *a*_0_ = 0 liftoff point (Figure 7 bottom panel, dashed vertical line). To understand this behavior, note that an instantaneous perturbation of *x*_*r*_ directly propagates to neural pools through sensory feedback. While all three neural pools are affected by this mechanical perturbation, the neural components of *η*^close^ are zero most of the time except when the trajectory lifts off from the *a*_0_ = 0 constraint (Figure 7 top panel, blue spike). This observation implies that the system has a high local sensitivity to *a*_0_ during the blue spike, whereas the sensitivity to *a*_1_ and *a*_2_ are significantly smaller than unity at all times. Thus, to understand the effect of perturbing *x*_*r*_ on the local timing, it is sufficient to focus on 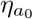 and examine how *a*_0_ reacts to perturbing *x*_*r*_.

Similar to the forward variational analysis, perturbing *x*_*r*_ delays the activation of *a*_0_, i.e., *a*_0_ lifts off from *a*_0_ = 0 at a later time. That is, the displacement in *a*_0_ near the *a*_0_ = 0 liftoff point is negative. Since 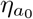 is negative near the liftoff point, perturbing *x*_*r*_ prolongs the total closed time (i.e., 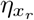 is positive during the closed phase).

Next we address the cusp phenomena observed in 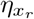 (Figure 7, bottom panel, yellow curve). Note that perturbations arriving before the trajectory lifts off from *a*_0_ = 0 delay the activation of *a*_0_ by increasing the inhibition from its sensory feedback. Moreover, the closer the time of perturbation to the time of liftoff, the larger the delay on the activation of *a*_0_. Such a larger delay leads to a greater increase of the total closed time due to perturbing *x*_*r*_. Hence, before the liftoff time (Figure 7, bottom panel, vertical black dashed line), 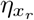 gradually increases. Once the trajectory has passed the liftoff point, perturbing *x*_*r*_ delays the activation of *a*_0_ by decreasing its sensory feedback, the effect of which now becomes excitatory. The size of this effect decays exponentially as the trajectory gradually leaves the boundary *a*_0_ = 0. Thus, there is a cusp in the 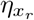 curve at the liftoff point, after which 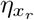 rapidly decreases.

### 2.4 Robustness to static perturbations

In this section, we show how the robustness of the *Aplysia* model (1.1), the ability of the system to maintain its performance despite perturbations, can be quantified using the ISRC, IPRC and LTRC analysis.

Following (Lyttle et al., 2017), we quantify the performance or task fitness via the average seaweed intake rate

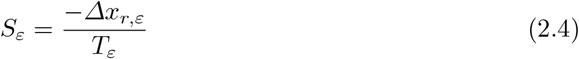

where *Δx*_*r,ε*_ is the net change in perturbed grasper position *x*_*r,ε*_ during the grasper-closed phase and *T*_*ε*_ is the perturbed period. Note that we assume the seaweed is moving together with the grasper when it is closed and not moving at all during the grasper-open component of the trajectory. Hence, −*Δx*_*r,ε*_ denotes the total amount of seaweed consumed per cycle.

Since the vector field *F*_*ε*_(**x**) in system (1.1) is piecewise smooth in the coordinates **x** and smooth in the perturbation *ε*, it follows that the following expansion holds:

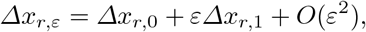

where *Δx*_*r*,0_ is the net change in the unperturbed grasper position during the grasper-closed component of the trajectory. Here, *Δx*_*r*,1_ is approximately given by the net change of the *x*_*r*_ component of the ISRC *γ*_1_, which is denoted as 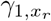 (see §2.2), over the grasper-closed phase. Suppose the grasper closes at *t*^close^ and opens at *t*^open^ over one cycle. It follows that 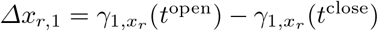.

(Lyttle et al., 2017) show that the robustness, i.e., the relative shift in the task fitness per relative change in perturbation, for small *ε*, can be written as

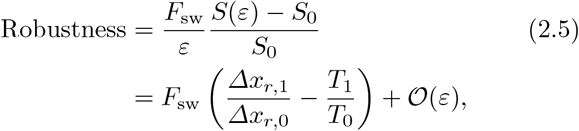

as *ε* → 0. Recall that *T*_0_ is the period of the unperturbed limit cycle and *T*_1_ denotes the linear shift in period, which can be estimated using the IPRC (see §2.3).

In summary, the robustness formula can be decomposed into two parts, one involving changes in shape (in particular, the grasper position *x*_*r*_) and the other involving the timing change. As discussed before, changes in shape can be estimated using the ISRC and the LTRC analysis, whereas the latter can be quantified using the IPRC. Below, we illustrate the quantification of the robustness by considering the perturbation to be the increase in the constant applied load *F*_sw_ → *F*_sw_ + *ε*.

The ISRC with or without timing rescaling corresponding to the perturbation on the applied load have already been computed and discussed in §2.1 and §2.2. Note that *Δx*_*r*,1_ in (2.5) is the net change in the ISRC during the grasper-closed phase. Choosing the ISRC with rescaling based on the timing of the closing and opening events provides a more accurate estimate of *Δx*_*r*,1_. Hence, we use the ISRC with piecewise rescaling to estimate *Δx*_*r*,1_, which is the net change in 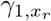 over the closing region per cycle (see the yellow curve over the shaded region in the lower right panel of Figure 5 and the green arrow marking the difference at the end of the closing phase). Furthermore, the linear shift in the period *T*_1_ can be estimated by (2.3) using the IPRC.

From analysis, we obtain 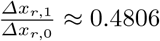 and 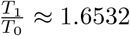, the above both of which are positive and are consistent with the concept of an adaptive “stronger and longer” change in the motor pattern in response to increased load. It follows that the robustness is approximately −1.1726 *×* 10^−2^. (Note that the smaller this number is in magnitude, the more robust the system is.) To the first order in *ε*, the relative change in the performance is then given by

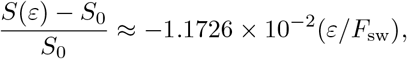

which is illustrated by the red circle as the perturbation size *ε* varies (see Figure 8, top panel). To see what this means, we take a data point on the line indicated by the arrow, i.e., (0.42, −0.005). Here *ε/F*_sw_ = 0.42 indicates a 42% increase in load *F*_sw_, which only causes a 0.5% decrease in the task fitness, corresponding to a highly robust response. Here the “stronger” effect (i.e., the first term in the robustness formula (2.5) being positive) contributes to the robustness whereas the “longer” effect (i.e., the second term in the robustness) reduces it. However, these two effects are not independent from each other: it is the longer retraction-closed time that allows the muscle to build up a stronger force, thereby contributing to a robust response.

**Fig. 8.**
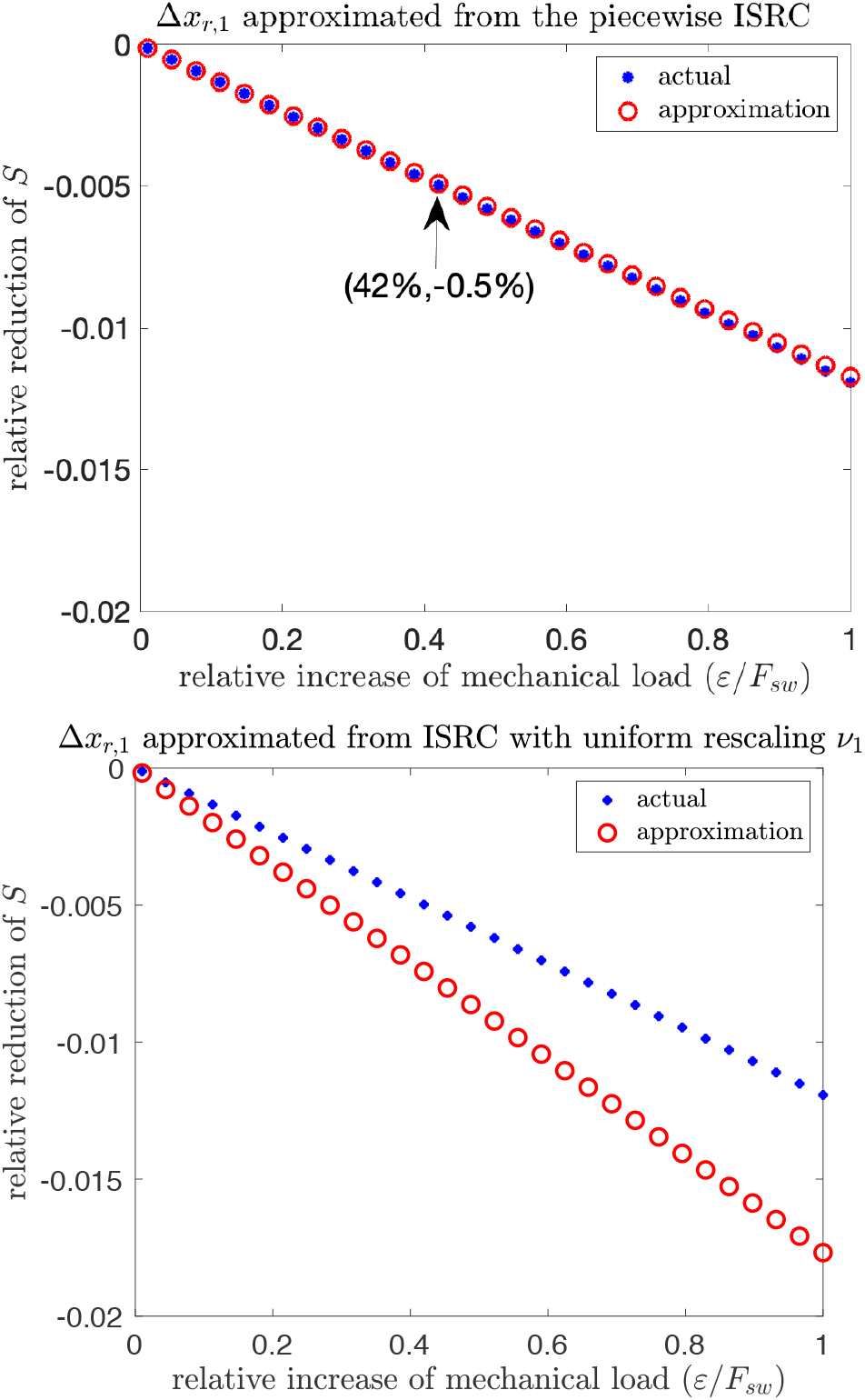
Relative change in task fitness (*S*(*ε*)−*S*_0_)*/S*_0_ computed numerically (blue stars) versus those obtained analytically from the ISRC and the IPRC according to formula (2.5) (red circles), as the perturbation *ε* on the seaweed load *F*_sw_ varies. Without perturbation, the nominal applied load is *F*_sw_ = 0.01. The approximation using the ISRC with different timing rescalings during the grasper-closed (*ν*_1,close_) versus grasper-open phases (*ν*_1,open_) estimated from the LTRC analysis matches the actual simulation (top panel), whereas the ISRC with uniform rescaling *ν*_1_ = *T*_1_*/T*_0_ estimated from the IPRC no longer gives a good approximation (bottom panel).

We also compute the relative change in *S* with respect to *ε* using direct numerical simulations (see Figure 8, blue stars), which show good agreement with our analytical results. In contrast, if we estimate *Δx*_*r*,1_ using the ISRC with a uniform timing rescaling (see Figure 11), the resulting estimated robustness becomes more negative and no longer gives an accurate approximation to the actual robustness (see Figure 8, bottom panel). That is, the ISRC using different rescaling factors over the grasper-closed phase (*ν*_1,close_) versus the grasper-open phase (*ν*_1,open_), gives a much better approximation to the robustness than the ISRC based on a global timing rescaling *ν*_1_ = *T*_1_*/T*_0_. The fact that the *ν*_1,open*/*close_ are obtained via the LTRC analysis highlights the contribution of this novel analytical tool. This observation demonstrates that for systems under certain circumstances (e.g, non-uniform perturbation as considered in system (1.1)), the ISRC together with the LTRC greatly improves the accuracy of the robustness, compared to the ISRC with global timing analysis given by the IPRC.

### 2.5 Sensitivity of robustness to other parameters

In general, the performance of motor control systems may be affected not only by external parameters, such as an applied load, but also be internal parameters, for instance describing the physical properties of the biomechanics or neural controllers. The variational tools used in the previous section to understand mechanisms of robustness to increases in applied load – the IPRC, ISRC and LTRC – can also give insights into the effects of changing internal model parameters. For instance, in the SLG model, appropriately varying strengths of protractor or retractor muscles can overcome effects of the increased mechanical load *F*_sw_ → *F*_sw_ + *ε*. Because of the SLG model’s relative simplicity, we can relate many of these changes to specific components of the fitness equation in detail.

Below, we first consider how varying sensory feedback strengths can help restore the reduced seaweed intake rate due to increased applied load. Then we examine how changing the strengths of the protractor and retractor muscles affects robustness to applied loads.

#### Varying sensory feedback strengths

Figure 9 shows the seaweed intake rate and robustness to the increased load *F*_sw_ with respect to changes in sensory feedback strengths *ε*_*i*_, *i* ∈ {0, 1, 2}. The performance *S*_0_ becomes negative when *ε*_0_ or *ε*_1_ is relatively small (e.g., smaller than 10^−5^) or when *ε*_2_ is relatively big (e.g., larger than 10^−3^), during which the system is in a fast limit cycle/biting mode and hence cannot swallow seaweed.

**Fig. 9.**
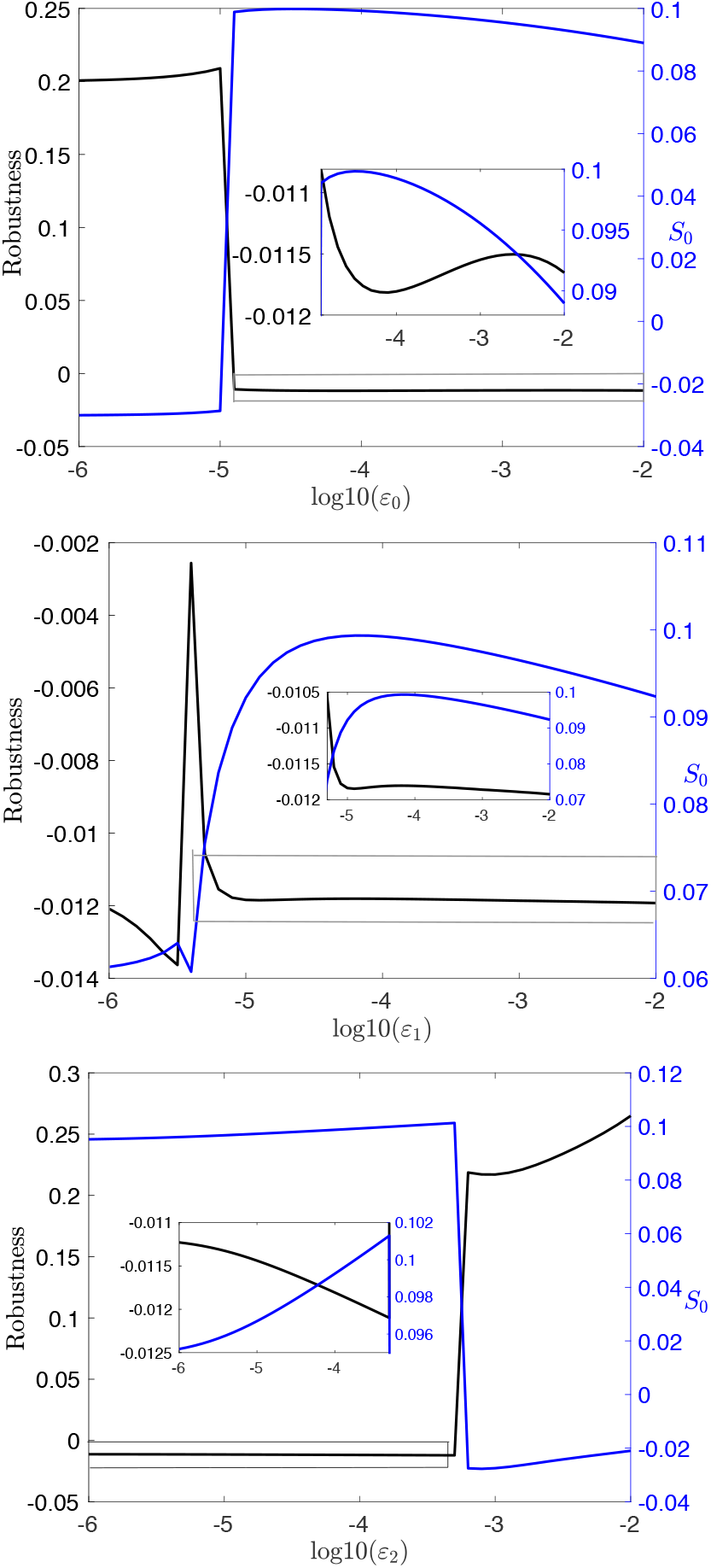
Effects of varying *ε*_0_ (top row), *ε*_1_ (center row) and *ε*_2_ (bottom row) on the robustness (2.5) to *F*_sw_ and the unperturbed seaweed intake rate *S*_0_, when *F*_sw_ = 0.01. Blue curves: Fitness *S*_0_. Black curves: Robustness.

When the system is in the heteroclinic/swallowing mode, as one might expect, increasing the sensory feedback (e.g., *ε*_2_) improves the performance. Surprisingly, our results show that increasing sensory feedback strengths to the two protraction-related neural pools leads to opposite results by decreasing the performance. These results seem to suggest that to restore the deficit caused by the increased load and achieve an increased robustness, we can either increase *ε*_2_ or decrease *ε*_0_ and/or *ε*_1_. However, this is not true. As shown in Figure 9, a decrease in the robustness can be induced by either decreasing *ε*_0_ or increasing *ε*_2_. Moreover, the robustness is largely insensitive to changes in *ε*_1_, despite the fact that it influences the performance. Understanding these effects on the robustness would require analysis of a second-order variational problem and represents a future direction for understanding neuromodulation.

#### Varying muscle strengths

Next we investigate how variations of *k*_0_ and |*k*_1_|, the strengths of the protraction and retraction muscles, affect the robustness to changes in seaweed load.

Figure 10 shows that performance improves with the increased protractor muscle strength *k*_0_ or the increased retractor muscle strength |*k*_1_|. This suggests that increasing *k*_0_ or |*k*_1_| can help restore the deficit in the performance due to the increased mechanical load and hence boost the robustness, which agrees with our numerical simulations (see Figure 10, top panel, black curve).

**Fig. 10.**
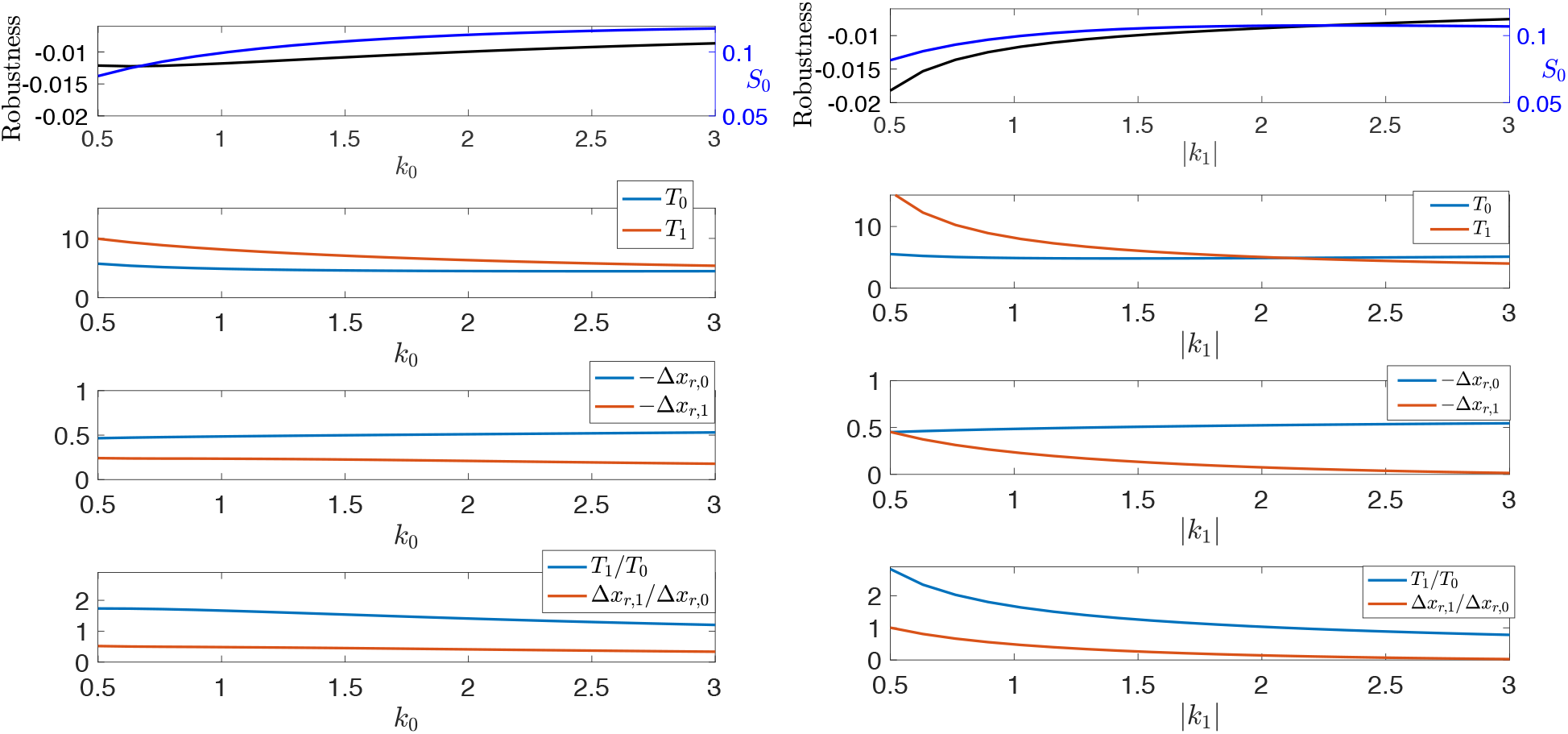
Effects of varying muscle strengths *k*_0_ (left panels) and |*k*_1_| (right panels) on the robustness to *F*_sw_ (top panels, black curve) and the unperturbed seaweed intake rate *S*_0_ (top panels, blue curve). Default parameters *k*_0_ = 1, *k*_1_ = −1 represent the strengths and directions of protraction and retraction muscles. The second and third rows of panels show the effects of muscle strengths on timing (*T*_0_, *T*_1_) and shape (−*Δx*_*r*,0_, −*Δx*_*r*,1_), respectively. The bottom panels shows how *T*_1_*/T*_0_ (blue) and *Δx*_*r*,1_*/Δx*_*r*,0_ (red) change as muscle strengths vary.

Recall that the robustness can be approximated as 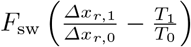 (see equation (2.5)). Understanding the underlying mechanisms of the robustness requires one to investigate how the two quantities involving shifts in shape and timing change with respect to *k*_0_ or |*k*_1_| (see Figure 10, lower three panels). We find that increasing *k*_0_ or |*k*_1_| reduces *T*_1_ and −*Δx*_*r*,1_ while *T*_0_ and −*Δx*_*r*,0_ are almost unaffected. Hence, both *Δx*_*r*,1_*/Δx*_*r*,0_ (the “stronger” effect in response to perturbations on the seaweed load) and *T*_1_*/T*_0_ (the “longer” effect) are decreased as we increase the muscle strengths. However, the reduction in the “stronger” effect is smaller than the reduction in the “longer” effect. As a result, the robustness approximated by 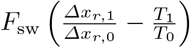 increases as *k*_0_ or |*k*_1_| increases.

Together, our analytical tools suggest ways in which coordinated changes in intrinsic parameters could maintain fitness and thus enhance robustness.

## 3 Methods

In this section, we review the classical variational theory for limit cycles (e.g., (Filippov, 1988; Bernardo et al., 2008; Leine and Nijmeijer, 2013; Park et al., 2018)), and new tools that we recently developed in Wang et al. (2021) for linear approximation of the effects of small sustained perturbations on the timing and shape of a limit cycle trajectory in both smooth and nonsmooth systems.

In the next two sections we treat the smooth and nonsmooth cases, respectively. In each case, we consider a one-parameter family of *n*-dimensional dynamical systems

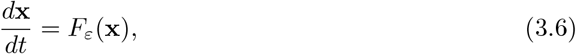

indexed by a parameter *ε* representing a static perturbation of a reference system

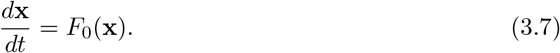

### 3.1 Timing and shape responses to static perturbations in smooth systems

Following Wang et al. (2021), we make the following assumptions:

#### Assumption 1

– *The vector field F*_*ε*_(**x**) : *Ω ×ℐ* → ℝ^*n*^ *is C*^1^ *in both the coordinates* **x** *in some open subset Ω* ⊂ ℝ^*n*^ *and the perturbation ε* ∈ *ℐ* ⊂ ℝ, *where ℐ is an open neighborhood of zero*.
– *For ε* ∈ *ℐ, system* (3.6) *has a linearly asymptotically stable limit cycle γ*_*ε*_(*t*), *with finite period T*_*ε*_ *depending (at least C*^1^*) on ε*.

It follows from Assumption 1 that when *ε* = 0, *F*_0_(**x**) is *C*^1^ in **x** ∈ *Ω* and the unperturbed system (3.7) exhibits a *T*_0_-periodic linearly asymptotically stable limit cycle solution *γ*_0_(*t*) = *γ*_0_(*t* + *T*_0_) with 0 *< T*_0_ *<* ∞. Assumption 1 also implies that the following approximations hold:

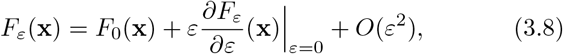

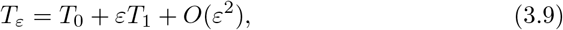

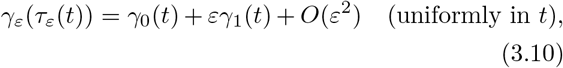

where *T*_1_ is the linear shift in the limit cycle period in response to the static perturbation of size *ε*. This global timing sensitivity, *T*_1_, is strictly positive if increasing *ε* increases the period. The perturbed time *τ*_*ε*_(*t*) satisfying *τ*_0_(*t*) ≡ *t* and *τ*_*ε*_(*t* + *T*_0_) − *τ*_*ε*_(*t*) = *T*_*ε*_ will be described later; it allows the approximation (3.10) to be uniform in time and permits us to compare perturbed and unperturbed trajectories at corresponding time points.

The timing and shape aspects of limit cycles are complementary, and may be studied together by considering the *infinitesimal phase response curve* (IPRC) and the *variational analysis* of the limit cycle, respectively.

#### Infinitesimal Phase Response Curve (IPRC)

The IPRC is a classical analytic tool that measures the timing response of an oscillator due to an infinitesimally small perturbation delivered at any given point on the limit cycle. It satisfies the adjoint equation (Schwemmer and Lewis, 2012)

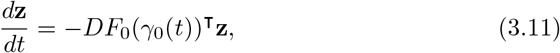

with the normalization condition

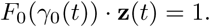

The linear shift in period *T*_1_ can be calculated using the IPRC as

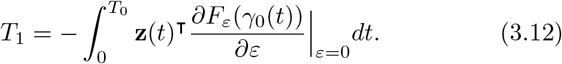

#### Forward Variational Equation

Classical sensitivity analysis (Wilkins et al., 2009) has been used in many applications to study the shape sensitivity or response of an oscillator to sustained perturbations.

The dynamics of the linear shift

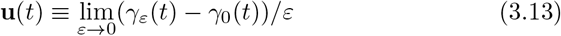

at time *t* of the periodic orbit *γ*_*ε*_(*t*) due to a sustained parametric perturbation *ε* initiated at time 0 satisfies the following forward variational equation:

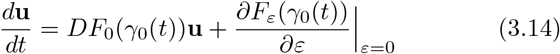

with initial condition **u**(0) set by the difference in the perturbed and unperturbed trajectories at the point where they cross the Poincaré section defined by the beginning of the closed phase. Specifically,

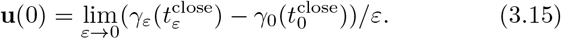

Compared with the homogeneous variational equation, which studies the shape sensitivity to instantaneous perturbations, the forward variational equation (3.14) contains a non-homogeneous term arising directly from the parametric perturbation acting on the vector field.

However, since the perturbed limit cycle has a different period *T*_*ε*_ and hence a different perturbed time *τ*_*ε*_ due to sustained perturbations, the forward variational equation which neglects such changes in timing fails to give a valid comparison between the perturbed and unperturbed trajectories for times on the order of a full cycle or longer (see Figure 2C and D). Hence, we adopt a new tool developed in Wang et al. (2021), the *infinitesimal shape response curve* (ISRC), which incorporates both the shape and timing aspects and captures a more accurate first-order approximation to the change in shape of the limit cycle under a parameteric perturbation.

#### Infinitesimal Shape Response Curve (ISRC)

Suppose the rescaled perturbed time can be written as *τ*_*ε*_(*t*) = *t/ν*_*ε*_ ∈ [0, *T*_*ε*_] for *t* ∈ [0, *T*_0_]. It follows that the relative change in timing denoted by *ν*_*ε*_ = *T*_0_*/T*_*ε*_ can be represented as *ν*_*ε*_ = 1 − *εν*_1_ + *O*(*ε*^2^) where 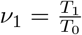.

Wang et al. (2021) denote the linear shift in the periodic orbit, *γ*_1_(*t*) in (3.10), as the ISRC and show it satisfies the following variational equation:

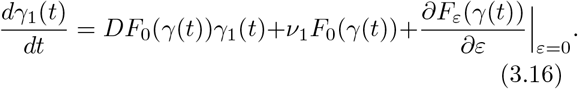

This equation resembles the forward variational equation (3.14), but has one additional non-homogeneous term arising from time rescaling *t* → *τ*_*ε*_(*t*). In contrast to the forward variational dynamics 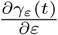, the ISRC *γ*_1_(*t*) is periodic with period *T*_0_ (see Figure 11, left). To see how well the ISRC approximates the actual linear shift between the perturbed and unperturbed trajectories, we plot the linear shift approximated from the ISRC (black curve) and the actual displacement (red dashed curve). Overall, they show good agreement with each other except near the transition between the grasper-closed and grasper-open phases. Such discrepancies arise from the fact that the solution segment at the closing phase has different timing sensitivity to the parametric perturbation compared with the segment at the opening phase, as discussed before. While these small errors are nearly unnoticeable (see Figure 11, right), they expand when the ISRC result is used to calculate the robustness (see Figure 8, bottom panel).

**Fig. 11.**
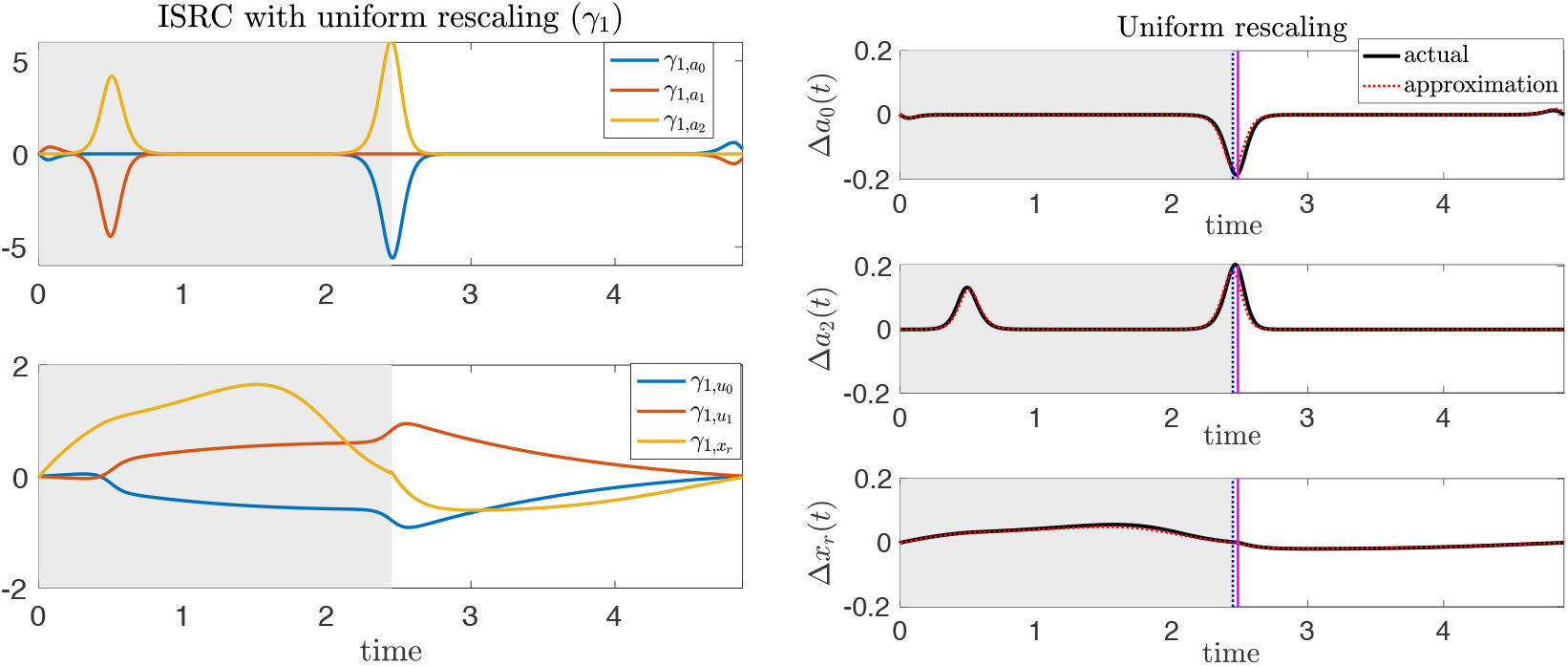
Variational analysis with uniform rescaling. The same perturbation as in Figure 2 is applied to the *Aplysia* model (1.1). Left: The ISRC *γ*_1_(*t*) with a uniform rescaling over one period. Right: Time series of the difference between the perturbed and unperturbed solutions along *a*_0_-, *a*_2_-, and *x*_*r*_-directions. The black curve denotes the numerical displacement (*Δy*(*t*) = *y*_*ε*_(*τ*_*ε*_(*t*)) − *y*(*t*)) computed by subtracting the unperturbed solution trajectory from the perturbed trajectory, after globally rescaling time, and aligning trajectories at the onset of closing. The red dashed curve denotes the product of the perturbation *ε* and the ISRC curve. The vertical blue dashed lines indicate the times at which the unperturbed grasper switches from closed to open. Shaded regions and the vertical magenta lines have the same meanings as in Figure 2. The perturbation is the same as in Figure 2.

Thus, in the case when a parametric perturbation leads to different timing sensitivities in different regions, we use the *local timing response curve* (LTRC) defined by Wang et al. (2021) to compute shifts in timing in different regions in order to improve the accuracy of the ISRC, as demonstrated when considering perturbations to the load applied to the seaweed (see Figure 12).

**Fig. 12.**
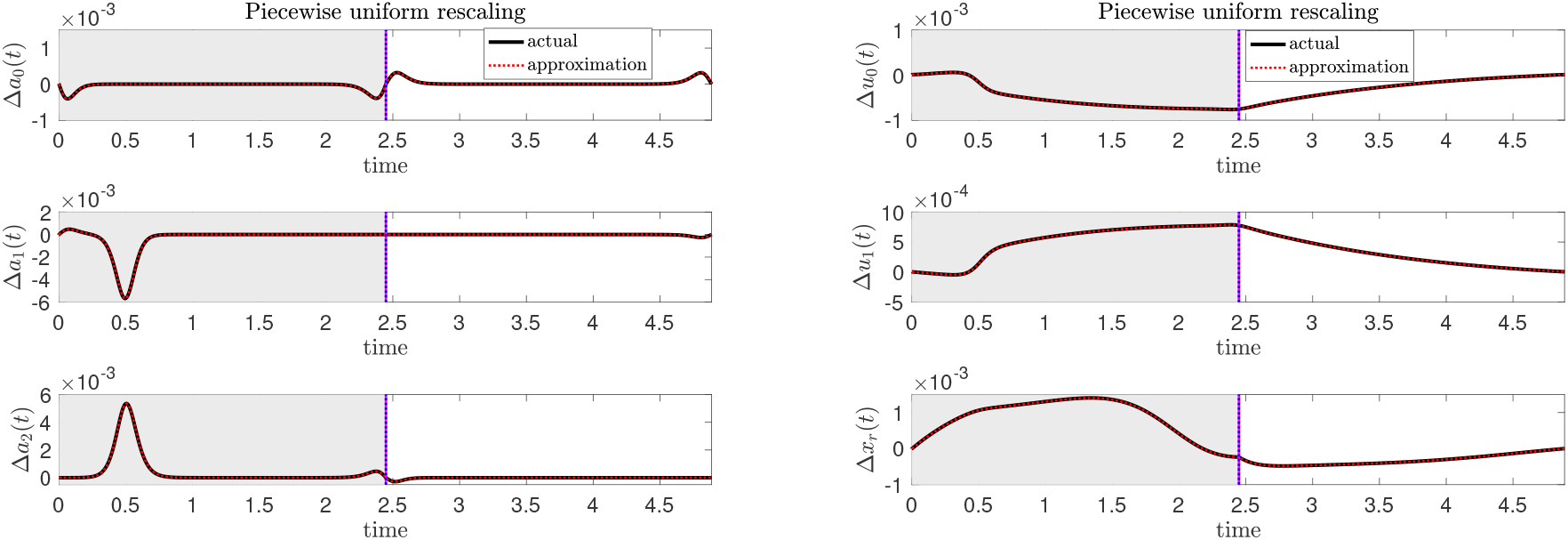
Displacements between perturbed and unperturbed trajectories estimated from the ISRC *γ*_1_ with piece-wise uniform rescaling (red dashed, *εγ*_1_) agree well with the actual displacement *Δy*(*t*) = *y*_*ε*_(*τ*_*ε*_(*t*)) − *y*(*t*) where *y* = {*a*_0_, *a*_1_, *a*_2_, *u*_0_, *u*_1_, *x*_*r*_} (solid black). Shaded regions, vertical magenta and blue lines have the same meanings as in Figure 11. The perturbation is the same as in Figure 2.

#### Local Timing Response Curve (LTRC)

The accuracy of the ISRC in approximating the linear change in the limit cycle shape evidently depends on its timing sensitivity, that is, the choice of the relative change in frequency *ν*_1_. In (3.16), we chose *ν*_1_ to be the relative change in the full period, by assuming the limit cycle has constant timing sensitivity. It is natural to expect that different choices of *ν*_1_ will be needed for systems with varying timing sensitivities along the limit cycle. To more accurately capture timing sensitivity of such systems to static perturbations, Wang et al. (2021) defined a *local timing response curve* (LTRC) which is analogous to the IPRC but measures the linear shift in the time that the trajectory spends within any given region. Specifically, the LTRC is the gradient of the time remaining in a given region until exiting it through some specified Poincaré section - a local timing surface corresponding to the exit boundary of this region. Such a section could be given as a boundary where the dynamics changes between regions, or where a perturbation is applied in one region but not another. For instance, in the feeding system of *Aplysia californica* (Shaw et al., 2015; Lyttle et al., 2017), the open-closed switching boundary of the grasper defines a local timing surface.

Let *η*^I^ denote the LTRC vector for region I. Suppose that at time *t*^in^, the trajectory *γ*_0_(*t*) enters region I upon crossing the surface *Σ*^in^ at the point **x**^in^; at time *t*^out^, *γ*_0_(*t*) exits region I upon crossing the surface *Σ*^out^ at the point **x**^out^. Similar to the IPRC, the LTRC *η*^I^ satisfies the adjoint equation

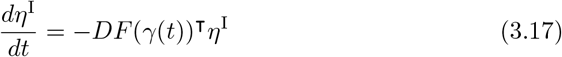

together with the boundary condition at the exit point

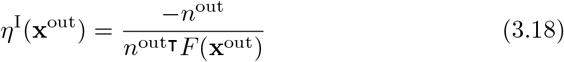

where *n*^out^ is a normal vector of *Σ*^out^ at the unperturbed exit point **x**^out^. The linear shift in the total time spent in region *I*, 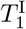, is given by

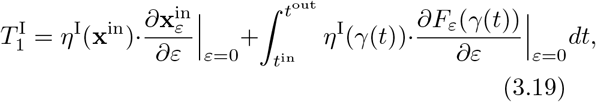

where 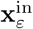 denotes the coordinate of the perturbed entry point into region I. It follows that the relative change in frequency local to region I is given by 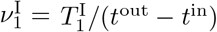.

#### Piecewise uniform ISRC

The existence of different timing sensitivities of *γ*(*t*) in different regions therefore leads to a piecewise-specified version of the ISRC (3.16) with period *T*_0_,

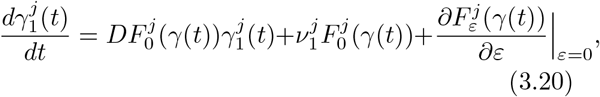

where 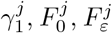 and 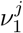 denote the ISRC, the unperturbed vector field, the perturbed vector field, and the relative change in frequency in region *j*, respectively, with *j* ∈ {I, II, III, *…*}. Note that in a smooth system, 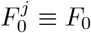 for all *j*.

As discussed before, the piecewise-specified ISRC, where *ν*_1_ takes different values in the closing and opening phases, nicely complements the forward variational analysis. It provides a more self-consistent global description of the shape response of the limit cycle to the mechanical perturbation (see Figure 5). Displacements between perturbed and unperturbed trajectories estimated using the piecewise-specified ISRC agree well with the actual displacements (see Figure 12). Moreover, it yields a much better approximation to the robustness compared with the ISRC with uniform rescaling (see Figure 8).

### 3.2 Timing and shape responses to static perturbations in nonsmooth systems

As discussed before, system (1.1) is a piecewise smooth system with one transversal crossing boundary *Σ*_*o/c*_ and three hard boundaries (*Σ*_0_, *Σ*_1_, *Σ*_2_). The study of limit cycle motions in such nonsmooth systems requires analytical tools beyond the standard arsenal of phase response curves and variational analysis, developed for systems with smooth (differentiable) right-hand sides (Spardy et al., 2011a,b; Park et al., 2017). For small instantaneous displacements, variational analysis has been extended to nonsmooth dynamics with both transversal crossing boundaries and hard boundaries for studying the linearized effect on the shape of a trajectory (Filippov, 1988; Bernardo et al., 2008; Leine and Nijmeijer, 2013; Dieci and Lopez, 2011). Analysis in terms of infinitesimal phase response curves (IPRC) has likewise been extended to nonsmooth dynamics for studying the linear shift in the timing of a trajectory following a small perturbation, provided the flow is always transverse to any switching surfaces at which nonsmooth transitions occur (Shirasaka et al., 2017; Park et al., 2018; Chartrand et al., 2018; Wilson, 2019). Recently, Wang et al. (2021) extended the IPRC method to nonsmooth systems with hard boundaries.

In nonsmooth systems with degree of smoothness one or higher (i.e., *Filippov systems*), the right-hand-side changes discontinuously as one or more switching surfaces are crossed. A trajectory reaching a switching surface or boundary has two behaviors: it may cross the boundary transversally or it may slide along it. Hence, there are two types of boundary crossing points: *transversal crossing points*, at which the trajectory crosses a boundary with finite velocity in the direction normal to the boundary, and *non-transversal crossing points* including the *landing point* at which a sliding motion along a switching boundary begins, and the *liftoff point* at which the sliding terminates. The time evolutions of the solutions to the variational equation (i.e., the forward variational dynamics and the ISRC) and the solutions to the adjoint equation (i.e., the IPRC and the LTRC) may experience discontinuities at a boundary crossing point (Filippov, 1988; Bernardo et al., 2008; Leine and Nijmeijer, 2013; Park et al., 2018; Wang et al., 2021).

The discontinuity in the variational dynamics when a trajectory meets a boundary crossing point **x**_*p*_ at crossing time *t*_*p*_ can be expressed with the saltation matrix *S*_*p*_ (see Table 1):

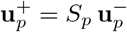

where **u**(*t*) denotes the solution of the forward variational equation or the ISRC, 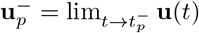 and 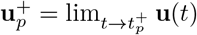 represent the solution just before and just after the crossing, respectively.

**Table 1.** Saltation matrices and jump matrices at boundary crossing points in Filippov systems (Filippov, 1988; Bernardo et al., 2008; Leine and Nijmeijer, 2013; Park et al., 2018; Wang et al., 2021)

|  | Landing Point | Transversal Crossing Point | Liftoff Point |
| --- | --- | --- | --- |
| Variational dynamics | $S_p = I - n_p n_p^\top$ | $S_p = I + \frac{(F_p^+ - F_p^-) n_p^\top}{n_p^\top F_p^-}$ | $S_p = I$ |
| IPRC & LTRC (forward time) | $J_p = I$ | $J_p = (S_p^{-1})^\top$ | $J_p$ is undefined |
| IPRC & LTRC (time-reversed) | $\mathcal{J}_p = I$ | $\mathcal{J}_p = J_p^{-1}$ | $\mathcal{J}_p = I - n_p n_p^\top$ |

In the table, $S_p$ , $J_p$ and $\mathcal{J}_p$ denote the saltation matrix, the jump matrix, and the time-reversed jump matrix at some boundary crossing point $\mathbf{x}_p = \mathbf{x}(t_p)$ , respectively. $F_p^- = \lim_{\mathbf{x} \rightarrow \mathbf{x}_p^-} F(\mathbf{x})$ and $F_p^+ = \lim_{\mathbf{x} \rightarrow \mathbf{x}_p^+} F(\mathbf{x})$ denote the vector fields of the nonsmooth system just before and just after the crossing at $\mathbf{x}_p$ , $I$ denotes the identity matrix, $n_p$ denotes the unit normal vector of the crossing boundary at $\mathbf{x}_p$ .

The discontinuity in **z**(*t*), the solution to the adjoint equation, at a boundary crossing point **x**_*p*_ may be expressed with the forward jump matrix (*J*_*p*_)

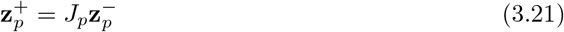

where 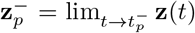 and 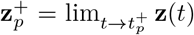 are the IPRC or the LTRC just before and just after crossing the switching boundary at time *t*_*p*_ in forwards time. However, Wang et al. (2021) showed that the jump matrix is not well defined at a liftoff point and hence introduced a time-reversed version of the jump matrix, denoted as *𝒥*_*p*_, defined as follows:

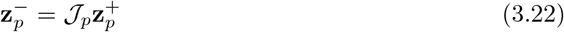

Table 1 summarizes the saltation and jump matrices at different types of boundary crossing points.

### 3.3 Simulation codes

Simulation codes written in Matlab are available at https://github.com/yangyang-wang/AplysiaModel.

## 4 Discussion

### Overview

Motor systems are robust - they maintain their performance despite perturbations. Understanding the mechanisms of robustness is important but challenging. To unravel the contributions of different components of robustness, we adopted tools we established in Wang et al. (2021) and reviewed in the methods section (§3) for studying combined shape and timing responses of both continuous and nonsmooth limit cycle systems under small sustained perturbations. We applied these tools to understand the mechanisms of robustness in a neuromechanical model of triphasic motor patterns in the feeding apparatus of *Aplysia* developed in (Shaw et al., 2015; Lyttle et al., 2017). We show in the results section (§2) that this framework lets us analyze how a small sustained perturbation alters the shape and timing of a closed loop system, and thus we began to describe how the neural and biomechanical components interact to contribute to robustness.

The first perturbation we considered was a sustained increase in mechanical load (*F*_sw_ → *F*_sw_ + *ε*). To our surprise, we discovered that long before sensory feedback affected the system, biomechanics played an essential role in robustness by producing an immediate force increase to resist the applied load (Figure 2, 3 and 4). Furthermore, although the sensory feedback immediately responded to the perturbation, its effect was delayed by the hard boundary properties of the neural firing rates. Our analysis suggests that sensory feedback contributes to the robustness primarily by shifting the timing of neural activation as opposed to changing neuronal firing rate amplitude (Figure 5, 6 and 7). Our methods can also be readily used to quantify how changes in timing and shape of trajectory affect the robustness (Figure 8). We find that sensory feedback and biomechanics contribute to the robustness of the system by generating a stronger retractor muscle force build-up during the prolonged retraction-closed phase that resists the increased load. The increased retractor muscle force ultimately leads to more seaweed being consumed during the slightly longer cycle time despite the large opposing forces, thereby contributing to a robust response. These new insights have refined and expanded a previous hypothesis that sensory feedback is the major mechanism that plays a crucial role in creating robust behavior (Lyttle et al., 2017).

Robustness is sensitive to other model parameters. For example, in §2.5 we investigated how varying internal parameters such as strengths of sensory feedback and muscle activity can help restore the performance that was reduced by an increased applied load (Figure 9 and 10). Again, we obtained some non-intuitive results. For example, increasing the sensory feedback strength can reduce the robustness rather than improving it (Figure 9). Moreover, increasing sensory feedback gain has opposite effects on performance and robustness, whereas increasing the protractor or retractor muscle strength improves both performance and robustness. Understanding sensitivities of performance to mixed parameters requires us to go beyond our existing methods. This second-order sensitivity represents an interesting future direction for understanding neuromodulation - the coordinated change of multiple system parameters in order to most effectively counter the effect of an external perturbation (Cropper et al., 2018). There are multiple pathways for neuromodulation, and the simplicity of the model lends itself to detailed analysis of multifactor sensitivities. In future work, we may apply the variational tools used in the present paper for understanding how changes in multiple parameters simultaneously could impact model performance and robustness (cf. §2.5).

### Experimentally testable predictions

The surprising result that the length-tension curves of the opposing muscles generate an instantaneous response to force perturbations could be tested, at least initially, using some of the more realistic biomechanics models that have been developed of *Aplysia* feeding.

For example, in a detailed kinetic model that does not have sensory feedback (Sutton et al., 2004; Novakovic et al., 2006), one could apply a step increase in force when the odontophore is closed and the retractor muscle is activated while measuring the force resistance to that change, and compare that to a purely passive response in which the retractor muscle is not activated. The results of this paper predict that there will be significant differences between these conditions.

In a model that does have sensory feedback (Webster-Wood et al., 2020), one could apply a step increase in force when the odontophore is closed and measure the change in force and the duration of the cycle to determine how that perturbation alters fitness. This paper’s results predict that the response to a sustained perturbation will be smaller in the presence of sensory feedback and will be larger if sensory feedback is removed.

The model suggests that there may be delays from the time that sensory feedback is available to the time that force changes. Using the model, sinusoidal force changes could be applied at different frequencies to determine the predicted phase lag, and this effect could be tested in the real animal.

Results shown in Figure 9 suggest that the model is relatively insensitive to changes in the strength of sensory feedback over a wide range of gains. Thus, one experimental test might be to increase or decrease the strength of sensory feedback to show that robustness to changing mechanical loads is not significantly affected. One way to test this hypothesis would be to use the newly developed technology of carbon fiber electrode arrays, which could be used to excite, inhibit, and record from many sensory neurons simultaneously (Huan et al., 2021).

In contrast, results shown in Figure 10 suggest that changing the relative strengths of the muscles can have larger effects on robustness. Previous studies have shown that neuromodulators can speed up and strengthen muscular contractions and thus might contribute to robustness (Taghert and Nitabach, 2012; Lu et al., 2015; Cropper et al., 2018). Studies of the neuromuscular transform (Brezina et al., 2000) suggested that neuromodulation could effectively speed up and strengthen feeding responses in normal animals, and thus might contribute to robustness.

Future experimental studies could be guided by coordinated changes of parameters in this model using the analysis tools we have presented.

### Caveats and limitations

Tracking possible transitions into and out of constraint surfaces becomes combinatorially complex as the number of distinct constraint surfaces grows. Here we impose three hard boundaries at *a*_*i*_ ≥ 0, as discussed above, by requiring firing rates to be nonnegative. An earlier model specification given in (Shaw et al., 2015; Lyttle et al., 2017) also required firing rates to be bounded via the constraint *a*_*i*_ ≤ 1. Here we relax this constraint for computational convenience, since the coexistence of multiple constraints requires encoding entry/exit conditions and vector field restrictions for all feasible combinations of constraints. In practice, comparison of simulations with and without the *a*_*i*_ ≤ 1 constraint give qualitatively and quantitatively indistinguishable results under most conditions.

Our analysis is in principle limited to small perturbations. Large perturbations lead to crossing of bifurcation boundaries in which the behavior switches to a different dynamical mode. “Robustness” in a broader sense can mean the distance to a basin of attraction of another dynamical attractor. For example, if the force is increased too much, the model will collapse into a stable fixed point with overextended protraction, while the animal will engage a different response to release or sever the seaweed to avoid damage to its feeding apparatus. This aspect is not captured in the variational approach. Nonlinear and bifurcation analysis could complement the present study and is ripe for investigation in future work.

In this paper we considered a specific perturbation, namely increasing the force opposing seaweed ingestion *F*_sw_ → *F*_sw_ + *ε*. Note that in this formulation, the perturbation parameter *ε* carries the same units (force) as *F*_sw_. Consequently, in order to use a unitless measure of robustness, the expression (2.5) includes a factor of *F*_sw_*/ε*. Also, in this formulation, the timing sensitivity *T*_1_ (shift in period *per increase in force*) and shape sensitivity *γ*_1_ (shift in limit cycle shape *per increase in force*) have units including reciprocal force. As an alternative formulation, which might facilitate comparison of robustness to perturbations across different modalities, one could rewrite the force perturbation as *F*_sw_ → *F*_sw_(1 + *ε*). In this case *ε* would represent a unitless measure of *relative* perturbation size. The subsequent variational, IPRC, ISRC and LTRC analysis would remain unchanged, except the resulting quantities *Z, T*_1_, *γ*_1_, and *η*_1_ would undergo a change in units, hence a multiplicative (fixed) change in scale. An advantage of specifying perturbations as a relative or unitless quantity would be that a similar analysis to that undertaken in this paper could be applied to other modalities in the same or system or across disparate systems.

### Generalizability to other systems

Although we focused in the present work on the robustness of the mean rate of seaweed intake with respect to increases in the force opposing ingestion, our analysis carries over to other objective functions (e.g. calories consumed per energy expenditure) as well as other perturbations (e.g. temperature, which may alter the speed of feeding in *Aplysia*). The variational approach to analyzing robustness should apply to any reasonable (e.g. smoothly differentiable) objective function and any parameter represented in the system, e.g. adjustments to changes in speed, steepness, or right-left asymmetry of walking movements on a (split) treadmill system (Frigon et al., 2013; Embry et al., 2018).

The present manuscript applies variational methods to understand the robustness in a specific *Aplysia* neuromechanical model (Lyttle et al., 2017). This model makes significant simplifications to the real feeding apparatus control system in order to gain mathematical tractability and analytical and biological insights. Nonetheless, the framework developed in (Wang et al., 2021) applies naturally to more elaborate dynamical models of *Aplysia* feeding such as (Webster-Wood et al., 2020) and models incorporating conductance-based network descriptions of the central pattern generator (Cataldo et al., 2006; Costa et al., 2020). Thus, what we have done here provides a framework for understanding neural control of motor behaviors like the one considered in this paper.

More broadly, motor control beyond the *Aplysia* feeding system is also amenable to the analysis of the sort developed in §3 (Wang et al., 2021). For example, the stability of bipedal walking movements remains a challenge in the field of mobile robotics Vukobratovic et al. (2012); Westervelt et al. (2018). Biologically inspired robotics continues to provide alternative approaches with greater robustness than conventional devices (Beer, 2009; Pfeifer et al., 2007; Beer et al., 1997; Goldsmith et al., 2019). The variational framework exhibited here applies to these systems as well (Fitzpatrick et al., 2020). In the context of any motor control model, the variational analysis we present here should allow analysis of robustness of any reasonable objective function with respect to any system parameter.

## 5 Acknowledgments

This work was supported in part by National Institutes of Health BRAIN Initiative grant RF1 NS118606-01 to HJC and PJT, by NSF grant DMS-2052109 to PJT, and by NSF grant DBI 2015317 to HJC, as part of the NSF/CIHR/DFG/FRQ/UKRI-MRC Next Generation Networks for Neuroscience Program. This work was supported in part by the Oberlin College Department of Mathematics. We thank Zhuojun Yu for providing a critical reading of the manuscript.

## A Tables for model parameters and initial conditions

Values for model parameters and initial conditions of state variables are given in Table 2 and Table 3.

## B Different timing sensitivities to muscle perturbations

Here we explain why increasing the protractor (resp., retractor) muscle activation during the early closing phase leads to a phase delay (resp., phase advance), whereas increasing the muscle activations during the late closing phase lead to the opposite effects (see Figure 6 in §2.3).

Early in the closing phase (i.e., the protraction-closed phase), increasing *u*_0_ leads to a phase delay. This effect occurs because with larger *u*_0_ force, *x*_*r*_ protracts more, which prolongs the inhibition to *a*_0_ through sensory feedback (feedback to *a*_0_ is inhibitory when *x*_*r*_ *>* 0.5). Hence *a*_0_ activates at a later time and the switch from closed to open is delayed, corresponding to a phase delay (see Figure 13A).

**Fig. 13.**
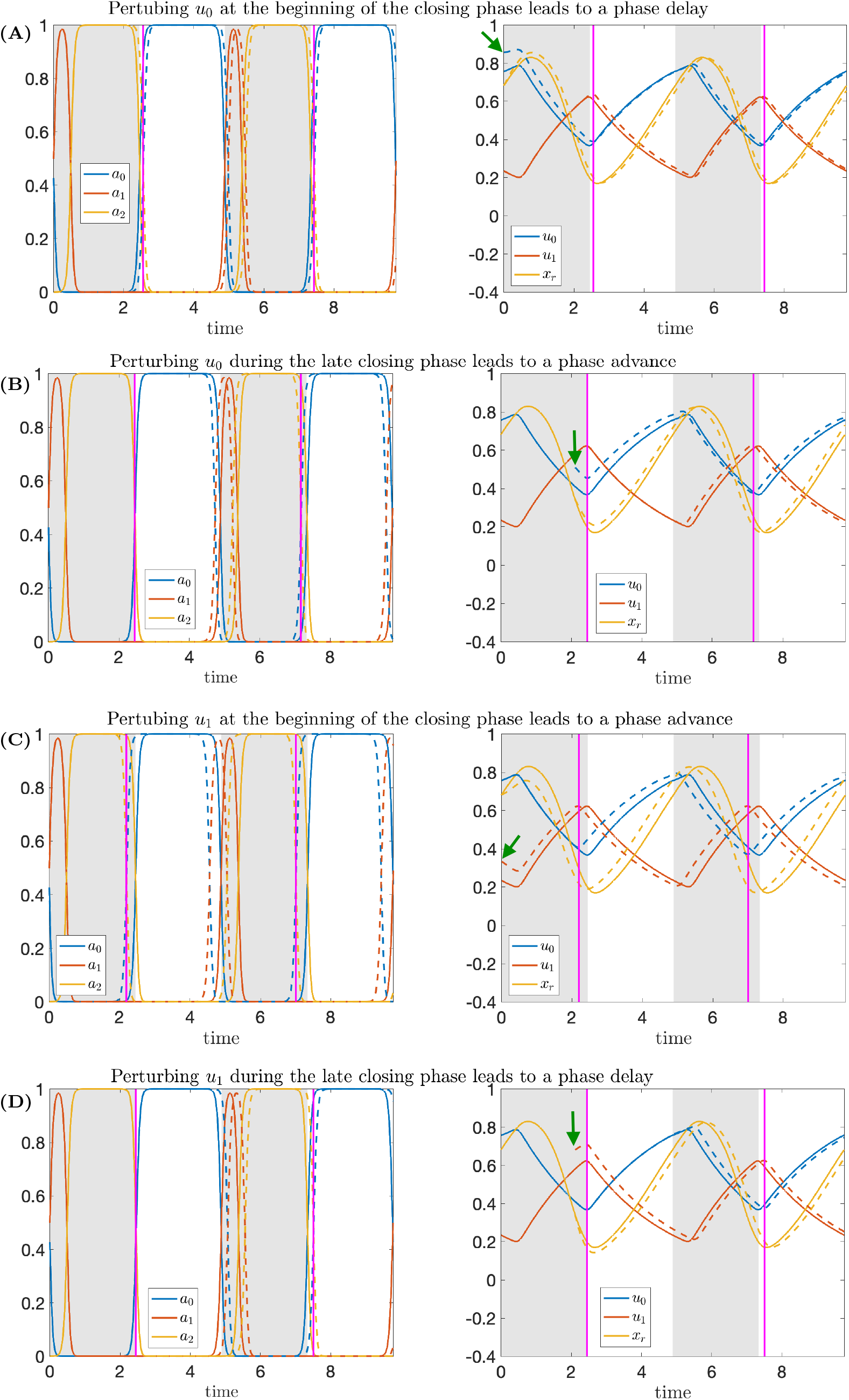
Time series of trajectories before (solid) and after (dashed) an instantaneous perturbation of the muscle activation variables (*u*_*i*_ → *u*_*i*_ + 0.1, see green arrows). Left panels show trajectories for neural variables, while right panels show trajectories for mechanical variables. (A) Perturbing the protractor muscle activation *u*_0_ at the beginning of the closing phase leads to a phase delay. (B) Perturbing *u*_0_ during the late closing phase leads to a phase advance. (C) Perturbing the retractor muscle activation *u*_1_ at the beginning of the closing phase leads to a phase advance. (D) Perturbing *u*_1_ during the late closing phase leads to a phase delay. Shaded regions and vertical magenta lines have the same meanings as in Figure 2.

On the other hand, increasing *u*_1_ during the early closing phase leads to a phase advance, because *x*_*r*_ decreases due to the increased retraction muscle forces and hence the inhibition switches to excitation earlier than in the original case (see Figure 13C).

During the late retraction-closed phase, increasing *u*_0_ leads to a phase advance (see Figure 13B). With increased protractor muscle force, *x*_*r*_ increases, but soon the state transitions to protraction-open. Then, the inhibition on *a*_1_ from the sensory feedback (feedback to *a*_1_ is inhibitory when *x*_*r*_ *<* 0.5) will be released earlier than before, because *x*_*r*_ is larger under perturbation and hence *a*_1_ activates earlier. As a result, the system switches from opening to closing phase earlier and this change corresponds to a phase advance.

On the other hand, if we increase *u*_1_ during the late closing phase, a phase delay results because *x*_*r*_ decreases with the perturbation. This effect prolongs the inhibition from sensory feedback to *a*_1_, since *x*_*r*_ stays below 0.5 for a longer time (see Figure 13D).

